# Cytoskeletal activation of NHE1 regulates mechanosensitive cell volume adaptation and proliferation

**DOI:** 10.1101/2023.08.31.555808

**Authors:** Qin Ni, Zhuoxu Ge, Yizeng Li, Gabriel Shatkin, Jinyu Fu, Kaustav Bera, Yuhang Yang, Yichen Wang, Anindya Sen, Yufei Wu, Ana Carina Nogueira Vasconcelos, Andrew P. Feinberg, Konstantinos Konstantopoulos, Sean X. Sun

## Abstract

Mammalian cells can rapidly respond to osmotic and hydrostatic pressure imbalances during an environmental change, generating large fluxes of water and ions that alter cell volume within minutes. While the role of ion pump and leak in cell volume regulation has been well-established, the potential contribution of the actomyosin cytoskeleton and its interplay with ion transporters is unclear. We discovered a cell volume regulation system that is controlled by cytoskeletal activation of ion transporters. After a hypotonic shock, normal-like cells (NIH-3T3, MCF-10A, and others) display a slow secondary volume increase (SVI) following the immediate regulatory volume decrease. We show that SVI is initiated by hypotonic stress induced Ca^2+^ influx through stretch activated channel Piezo1, which subsequently triggers actomyosin remodeling. The actomyosin network further activates NHE1 through their synergistic linker ezrin, inducing SVI after the initial volume recovery. We find that SVI is absent in cancer cell lines such as HT1080 and MDA-MB-231, where volume regulation is dominated by intrinsic response of ion transporters. A similar cytoskeletal activation of NHE1 can also be achieved by mechanical stretching. On compliant substrates where cytoskeletal contractility is attenuated, SVI generation is abolished. Moreover, cytoskeletal activation of NHE1 during SVI triggers nuclear deformation, leading to a significant, immediate transcriptomic change in 3T3 cells, a phenomenon that is again absent in HT1080 cells. While hypotonic shock hinders ERK-dependent cell growth, cells deficient in SVI are unresponsive to such inhibitory effects. Overall, our findings reveal the critical role of Ca^2+^ and actomyosin-mediated mechanosensation in the regulation of ion transport, cell volume, transcriptomics, and cell proliferation.

## Introduction

Cells actively control their size during the cell cycle, and failure of cell size control is an indicator of disease (*1–5*). Cell size or volume influences all intracellular biochemical reactions by affecting concentrations of cytoplasmic components. During interphase, the cell size steadily increases due to the synthesis of new cytoplasmic components (*6–9*). On the other hand, mitotic cells can swell and increase their volume by 10-20% in minutes without producing extra mass (*10, 11*). Moreover, short timescale cell swelling/shrinkage can generate significant mechanical forces (*10, 12–14*), provide cytoplasmic space for chromosome segregation (*15*), and also impact cell migration (*16–19*).

At the timescale of minutes, cell volume changes are predominantly due to water flux across the cell membrane, driven by both osmotic and hydraulic pressure gradients (*13, 18, 20*). When the cell size and shape are steady, hydraulic pressure gradients are also mechanically balanced by tension in the cell membrane and the actomyosin cortex (*14*). As a consequence, cell volume is sensitive to both environmental physical and chemical variables, including extracellular osmolarity (*1,18,21,22*), media viscosity (*19,23*), substrate stiffness or geometry (*24–27*), hydraulic resistance (*19, 28, 29*), and mechanical confinement (*30*).

To maintain the necessary balance between the osmotic pressure and the hydraulic pressure, cells use transmembrane ion fluxes to change the cytoplasmic solute concentration. Ion “pump and leak” has been identified as a major mechanism in volume regulation (*1, 31*), involving Na^+^ and K^+^ fluxes through ion pumps, exchangers and channels such as sodium-potassium ATPases (NKA) and sodium-hydrogen exchanger (NHE). The latter has been shown to be involved in mitotic swelling and cell migration (*11, 16, 19*). Actomyosin contractile force has been proposed as another way to control cell volume (*12, 13, 32–35*). However, direct evidence showing volume regulation by actomyosin is lacking. A central question that arises is **whether actomyosin directly influences cell volume via contractile force or indirectly regulates cell volume via signals to ion transporters.** Ca^2+^ related signaling cascades initiated by mechanosensitive ion channels is a promising candidate that bridge actomyosin contractility with ion transport (*1, 36–38*). Currently, how cell volume regulation is achieved through actomyosin and Ca^2+^ mediated signaling is still unclear.

In this work, we investigate the interplay between actomyosin dynamics, Ca^2+^ signaling, and ion fluxes in regulating cell volume. Our findings reveal that different cell types display distinct volume adaptation dynamics during hypotonic stress. We identify a secondary volume increase (SVI) in many normal-like cell lines, such as NIH-3T3 fibroblasts and non-tumorigenic breast MCF-10A cells, after the initial regulatory volume decrease (RVD). The SVI is triggered by cytoskeletal activation of NHE1 via Ca^2+^ influx through the mechanosensitive channel Piezo1, actomyosin remodeling, and the NHE1-actin binding partner ezrin. The cytoskeletal activation of NHE1 is also validated by direct mechanical stretching of cells. This mechanism appears to be mostly absent in certain cancer cell types such as HT1080 fibrosarcoma and MDA-MB-231 metastatic breast cancer cells. Importantly, 3T3 cells lack SVI when plated on compliant substrates, further demonstrating the sensory role of actomyosin in this process. Further explorations reveal that SVI leads to large-scale nuclear deformation and significant changes in the cell transcriptomic and epigenetic landscapes. Specifically, SVI uniquely up-regulates phosphatases that inhibit the ERK/MAPK signaling pathway, thereby impeding cell growth - a phenomenon that is not observed in cells lacking SVI.

## Results

### Hypotonic shock triggers non-monotonic cell volume responses

To reveal the cell volume regulatory system, we applied 50% hypotonic shock as an external perturbation and monitored the subsequent cell volume dynamics using the single-cell tracking fluorescence exclusion method (FXm, Fig. 1a) (*39, 40*). We first examined HT1080 fibrosarcoma cells and found that these cells underwent monotonic RVD within 20 min after hypotonic shock, reaching a steady state that is 10% higher than their isotonic volume (Fig. 1b, c). Hypotonic stress also resulted in a slight increase in HT1080 base area without significant cell shape change (Fig. S1a, b, and Video 1). Similar monotonic RVD with a faster recovery speed was found in MDA-MB-231 breast cancer cells (Fig. 1d) and also recently reported for Hela Kyoto cells (*21, 22*). The RVD showcases the ability of cells to quickly adjust their volume in response to environmental stimuli, and has been established as a universal property of mammalian cells (*1, 20*).

**Figure 1:**
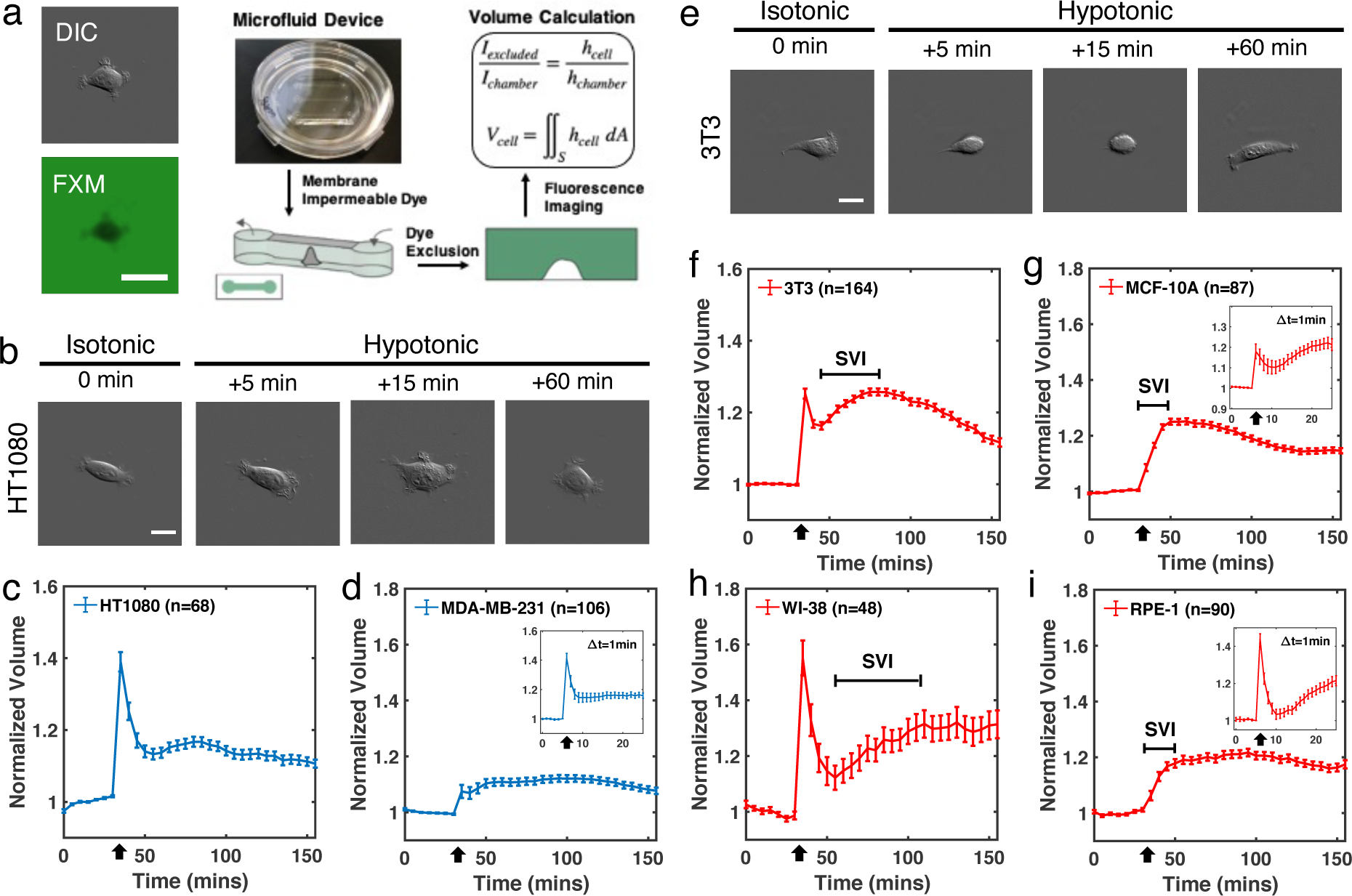
Distinct cell volume dynamics during hypotonic shock. (a) A schematic of the Fluorescence Exclusion method (FXm), and representative differential interference contrast (DIC) and epifluorescence images. (b) DIC images of HT1080 before and 5, 15, and 60 min after hypotonic shock. (c, d) Cell volume tracking of HT1080 (c), MDA-MB-231 (d) before and after hypotonic shock. Hypotonic solutions were injected into FXm channels at 30 min. Insets in (d) are volume imaged at 1 min frame rate for MDA-MB-231 (n=35). (e) Representative DIC images of 3T3 before and 5, 15, and 60 min after hypotonic shock. (f-g) Cell volume tracking of 3T3 (f), MCF-10A (g), WI-38 (h), and RPE-1 (i) before and after hypotonic shock. Insets in (g) and (i) are volume imaged at 1 min frame rate for MCF-10A (n=29) and RPE-1 (n=55), respectively. (c,d, f, g) Error bars represent the standard error of mean (SEM). Black arrows indicate the time of osmotic shock. (a, b, e) Scale bars, 20 µm.

Interestingly, normal-like 3T3 fibroblast cells during hypotonic shock displayed a second phase of volume increase after the initial RVD (Fig. 1e, f). 3T3 volume first decreased to a level that *∼*18% higher than its baseline isotonic volume after 10 min of hypotonic shock, accompanied with *∼* 30% base-area reduction (Fig. S1a) and significant cell rounding (Fig. S1b, c, and Video 2). After that, 3T3 showed a secondary volume increase (SVI), followed by another phase of volume decrease and re-spreading with increased base area. This SVI was also found in many other normal-like cell lines, including MCF-10A breast epithelial cells (Fig. 1g), WI-38 lung fibroblast cells (Fig. 1h), hTERT-immortalized retinal pigmented epithelial (RPE-1) cells (Fig. 1i), human embryonic kidney (HEK) 239A cells (Fig. S1d), and human foreskin fibroblast (HFF-1) cells (Fig. S1e), indicating that the SVI is common in many cell types and not a 3T3 specific response. Tracking single cell volume trajectories revealed significant cell to cell variations, both SVI and the monotonic RVD dynamics were found in all cell lines examined (Fig. S1f,g). The majority of 3T3 and MCF-10A cells displayed the SVI phenotype with varying timescales (on average 45 min for 3T3 and 42 min for MCF-10A, Fig. S1h) and magnitudes (on average 18% for 3T3 and 30% for MCF-10A, Fig. S1i). Since the doubling time of 3T3 is *∼* 17 h, SVI is not a consequence of cell growth. On the other hand, 60% hypertonic shock resulted in rapid cell volume decrease without significant regulatory volume increase in both 3T3 and HT1080 (Fig. S1j, k).

### Ion pump and leak regulates RVD and SVI

Pump and leak through ion transporters has been implicated as a key mechanism of cell volume regulation (*1, 31*), and thus we examined its impact on SVI. We first investigated the role of NHE, whose activation leads to an increase of both cell volume and intracellular pH by importing Na^+^ and exporting H^+^ (*41–43*). In isotonic conditions, adding the NHE inhibitor EIPA reduced cell volume in 3T3 but not in HT1080 (Fig. 2a). During hypotonic shock, HT1080 and MDA-MB-231 cells treated with EIPA displayed a significantly faster RVD (Fig. 2b, and Fig. S2c). This indicates that hypotonicity triggers NHE activation, leading to cell volume increase that impedes RVD. In 3T3 and MCF-10A cells, EIPA not only increased the speed of RVD, but also completely removed the SVI (Fig. 2c and Fig. S2d). Specifically, we found that the removal of SVI was controlled by NHE isoform 1 (NHE1) as confirmed by NHE1 knockdown using RNA interference (shNHE1, Fig. 2d and Fig. S2f), and the selective NHE1 inhibitor Bix (Fig. S2e). We then used an intracellular pH indicator, pHrodo Red-AM (*19, 44*), to assess NHE activity. Upon hypotonic shock, 3T3 displayed significant alkalinization that slowly recovered over 40 min, in striking contrast to the immediate decrease and rapid pH recovery found in HT1080 (Fig. 2e). These results confirm that NHE1 is activated after hypotonic shock and its activity is sustained during SVI in 3T3, whereas NHE1 activation in HT1080 occurs transiently during RVD. Overall, our observations indicate that NHE plays a significant negative role in RVD, and the observed SVI is associated with NHE1 activity.

**Figure 2:**
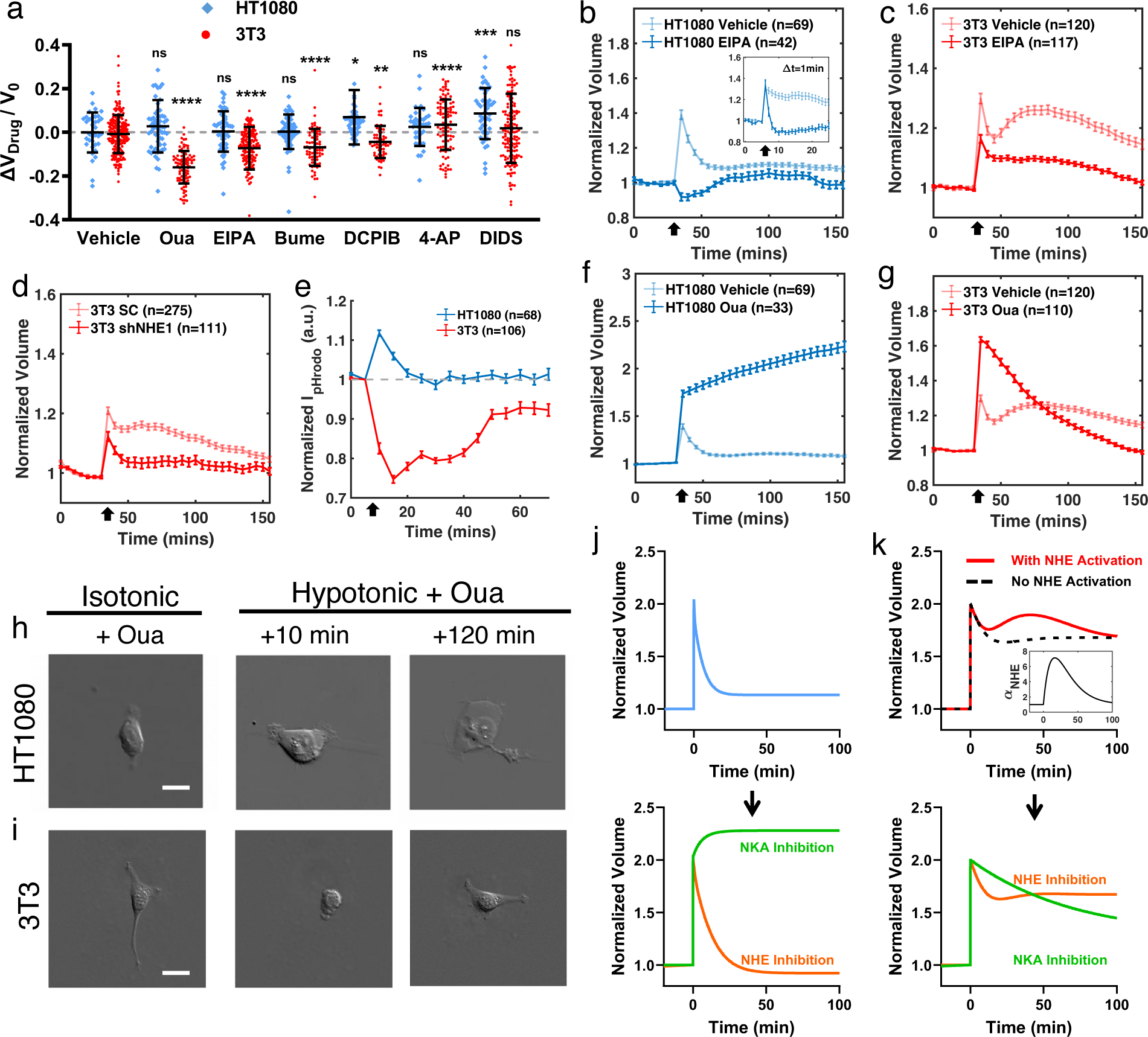
Ion fluxes regulate cell volume dynamics after hypotonic shock. (a) Fractional cell volume change after exposing to ion transporter inhibitors: Ouabain (Oua, NKA inhibitor), EIPA (NHE inhibitor), Bumetanide (Bume, NCKK inhibitor), DCPIB (VRAC inhibitor), and 4-AP (VGKC inhibitor). Cell volumes were tracked for 1-4 h until fully adapted (see methods for details). n_HT1080_= 36, 60, 50, 61, 47, 36, 47 and n_3T3_ = 213, 94, 147, 76, 62, 100, 147, respectively. (b, c) EIPA treated HT1080 (b) and 3T3 (c) volume dynamics under hypotonic shock. Inset in (b) has 1 min frame rate (Vehicle n = 53, EIPA n = 25). (d) 3T3 scramble control (SC) and shNHE1 volume dynamics under hypotonic shock. (e) Intensity of pHrodo-red AM for HT1080 and 3T3. Hypotonic shock was applied at 5 min. An increase in pHrodo-red intensity indicates decrease in pH. (f, g) Oua treated HT1080 (f) and 3T3 (g) volume under hypotonic shock. (h, i) Representative DIC images of Oua treated HT1080 (h) and 3T3 (i) before, 10 min, and 120 min after hypotonic shock. (j, k) Computer simulation of HT1080 (j) and 3T3 (k) with hypotonic shock and inhibition of NKA and NHE. Inset in (k) indicates the NHE activation function, where *α*_NHE_ is the permeability coefficient of NHE. See Supplementary Text for model details. (a) Error bars indicate standard deviation. Kruskal–Wallis tests followed by Dunn’s multiple-comparison test were conducted between data sets and the vehicle control of the corresponding cell type. (b-g) Error bars indicate the SEM. Mann–Whitney U-tests for two-condition comparisons were conducted at each time point post-shock, and p values were plotted as a function of time in Supplementary Text 3. (h,i) Scale bars, 20 µm.

We next examined NKA, which regulates cell volume decrease by importing 2 K^+^ and exporting 3 Na^+^ (*45*), using its inhibitor Ouabain (Oua). With high dosage (250 µM) of Oua, 3T3 volume decreased by 20% while HT1080 volume remained the same in isotonic media (Fig. 2a). During hypotonic shock, however, Oua completely removed HT1080’s RVD, where its initial cell volume peak (defined as the first measured time point after hypotonic shock) reached 60% higher than its isotonic volume (Fig. 2f). HT1080 volume continuously increased over the next 2 h, leading to significant cell swelling (Fig. 2h, Video 3). Similar results were observed in MDA-MB-231 and MCF-10A (Fig. S2a, b), confirming that NKA is a major contributor to RVD. Blocking NKA also increased the initial cell volume peak of 3T3 cells, but their volume still recovered (Fig. 2g, i). These markedly different behaviors suggest completely different cell volume regulatory mechanisms exist in tissue cells.

We also examined the effects of other key ion transporters in volume regulation, chosen based on their high mRNA levels in both cell lines (Supplementary Text-1), and their established role in cell volume regulation (*1*). In both 3T3 and HT1080, inhibition of Na^+^-K^+^-Cl*^−^*cotransporter (NKCC) using bumetanide (bume, Fig. S2g, h), volume-regulated anion channel (VRAC) using DCPIB (Fig. S2i, j), or Cl*^−^* and HCO*^−^* transporters using DIDS (Fig. S2m, n) had limited impact on the volume dynamics, while blocking passive voltage-gated K^+^channel (VGKC) using 4-Aminopyridine (4-AP) significantly increased the steady state volume after hypotonic shock (Fig. S2k, l). However, none of these inhibitors generated volume change more than 10% in isotonic environment (Fig. 2a), or changed the features of the monotonic RVD in HT1080 and the SVI in 3T3 during hypotonic shock.

### Theoretical modeling can explain cell type specific volume regulatory behaviors

To probe the potential mechanism underlying the distinct cell volume regulation observed between 3T3 cells and HT1080 cells, we employed a computational cell volume model (*42*) that considers fluxes from commonly expressed ion transporters, H^+^ and HCO*^−^*, and membrane electrical potential. In the model, the cell volume is modulated by osmotic and hydrostatic pressure gradients. Ion fluxes are determined by the activities of ion transporters (related to regulation of channel opening and their expression levels) as well as the chemical and mechano-physiological states such as cell cortical tension, pH, and membrane potential. The description of the model can be found in Supplementary Text-2. HT1080 and 3T3 cells display distinct volume adaptation dynamics, both immediately after osmotic change and at longer time scales over several minutes. Therefore, we looked for different parameter regimes in the model that reproduce these distinct behaviors. Using an unbiased search of ion transporter activity coefficients and conducting a principal component analysis (PCA), we indeed found two regions in parameter space that correspond to the types of cell volume regulation behavior observed in HT1080 and 3T3 cells (Fig. 2j, k, and Supplementary Text-2). PCA confirmed that NHE and NKA are two key regulators in initial RVD, in agreement with the experimental observations. The two sets of parameters also correctly predict the slightly higher homeostatic cytoplasmic pH observed in HT1080 versus 3T3 (Fig. S2p). On the other hand, experimental data suggests that NHE activation at longer time scales of tens of minutes generates SVI in 3T3. Our modeling results show that such complex secondary volume dynamics can only be achieved with additional NHE activation (Fig. 2k). By incorporating a second phase of NHE activation into its activity coefficient, the model can generate the same type of volume increase as the SVI in 3T3 (Fig. S2o). These findings raise the question of how this activation is regulated in different cell types, and what is the purpose of SVI.

### Actomyosin is involved in activating SVI

Hypotonic shock not only alters intracellular osmolarity, but also exerts forces on the actomyosin network and the cell membrane by expanding the cell surface (*13*). As a consequence, hypotonic stress has been reported to increase cell membrane tension (*21*) and lead to actin network reorganization (*46*). Thus, we examined the F-actin dynamics after hypotonic shock in 3T3 and HT1080 with EGFP-F-tractin. Confocal microscopy showed that in 3T3 cells, F-actin remodeled rapidly and formed a rounded actin cortex after osmotic shock (Fig. 3a), accompanied by elevated membrane tension measured by membrane tension probe Flipper-TR (*21, 47*) and Fluorescence Lifetime Imaging Microscopy (Fig. 3d, e). The cortex reorganization and high membrane tension were sustained during SVI. Quantitative immunofluorescence (IF) of phosphorylated myosin light chain (pMLC) also showed that 3T3 pMLC content increased after hypotonic shock and recovered after 60 min (Fig. S3a). In contrast, hypotonic stress neither triggered actin network remodeling in HT1080 (Fig. 3b) nor induced significant changes in the cell pMLC content (Fig. S3b). These observations suggest that 3T3 and HT1080 have distinct involvement of actomyosin machinery when responding to hypotonic stress, and we hypothesize that such difference is related to their distinct volume dynamics.

**Figure 3:**
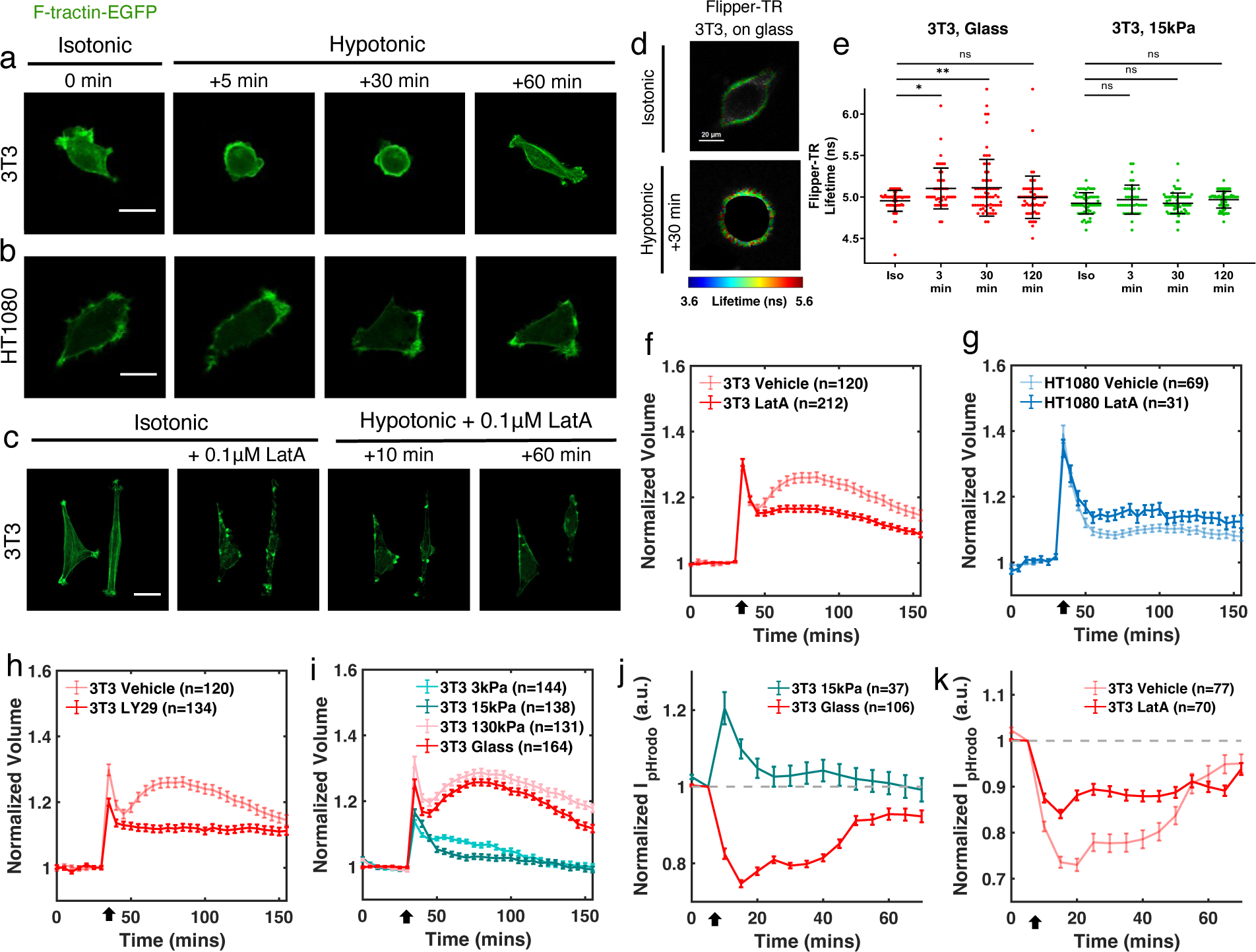
Actomyosin dynamics during hypotonic shock and its activation of SVI. (a-c) Confocal images of EGFP-F-tractin transfected 3T3 (a), HT1080 (b), and LatA treated 3T3 cells (c) before and after hypotonic shock. (d) Representative images showing elevated membrane tension measured by Flipper-TR lifetime during hypotonic shock in 3T3 cells grown on glass. (e) Membrane tension of 3T3 grown on glass and on 15 kPa PDMS substrates in before and after 3 min, 30 min, or 2 h hypotonic shock. n=61, 41, 72, 81 for glass, and 25, 18, 27, 40 for 15kPa substrates, respectively. (f-h) Volume dynamics of LatA treated 3T3 (f), LatA treated HT1080 (g), and LY29 treated 3T3 (h) under hypotonic shock. (i) Volume dynamics of 3T3 cells grown on 3, 15, and 130 kPa PDMS substrates and on glass under hypotonic shock. (j) Intensity of pHrodo-red AM for 3T3 grown on 15 kPa substrates versus on glass. Hypotonic shock was applied at 5 min. (k) Intensity of pHrodo-red AM for LatA treated 3T3 versus vehicle control under hypotonic shock. (f-k) Error bars indicate SEM. Mann–Whitney U-tests for two-condition comparisons were conducted at each time point post-shock, and p values were plotted as a function of time in Supplementary Text 3. (e) Error bars indicate standard deviation. Kruskal–Wallis tests followed by Dunn’s multiple-comparison test were used. (a-d) Scale bars, 20 µm.

We then examined whether the observed actomyosin remodeling influences cell volume. Addition of 100 nM actin depolymerizer Latrunculin A (LatA) was sufficient to disrupt the actin network and halt its remodeling after hypotonic shock in 3T3 cells (Fig. 3c). While LatA or another actin depolymerizer Cytochalasin D (CytoD) had limited impact on the initial RVD of 3T3 (Fig. 3f and Fig. S3c), myosin II inhibitor (S)-nitro-Blebbistatin (n-Bleb) reduced the initial peak (Fig. S3d). Interestingly, all three drugs removed SVI in 3T3. In addition to directly disrupting actomyosin, we inhibited phosphoinositide 3-kinases (PI3K) with LY294002 (LY29), known for activating actin protrusions (*48*). This treatment similarly reduced the initial peak and abolished SVI in 3T3 (Fig. 3h). We also validated the effect of LatA in removing SVI in MCF-10A (Fig. S3e). On the other hand, HT1080 cells treated with LatA still showed limited actin remodeling (Fig. S4a), and neither disrupting actomyosin using LatA, CytoD, and n-Bleb, nor inhibiting PI3K significantly affected monotonic RVD compared to untreated cells. (Fig. 3g, Fig. S3f-h).

Substrate stiffness is known to alter F-actin network structure and change cytoskeletal forces (*49–52*). As an independent way to validate the role of actomyosin in SVI, we cultured 3T3 cells on polydimethylsiloxane (PDMS) substrates with varying stiffness and examined cell volume dynamics during hypotonic shock. SVI was observed on both glass surfaces and stiff (130 kPa) substrates. In marked contrast and in line with our hypothesis, 3T3 displayed a monotonic RVD on soft substrates at 3 and 15 kPa (Fig. 3i). In addition, hypotonic shock led to transient pH reduction (Fig. 3j) and no detectable change of membrane tension (Fig. 3e) for 3T3 cells grown on 15 kPa substrates. On the other hand, HT1080 displayed consistent monotonic volume regulatory decrease regardless of substrate stiffness (Fig. S4b). Collectively, we conclude that SVI generation requires cytoskeletal activity and cellular mechanosensation of the mechanical microenvironment.

Our results demonstrate that SVI requires an active and intact actomyosin network. However, the lack of SVI after actomyosin disruption appears counterintuitive. Disruption of actomyosin reduces the contractile stress and thus, from the perspective of force balance, should have resulted in a volume increase (*12, 13*). Contrary to this, pharmacological inhibitions aimed at various components of the actin cytoskeleton or its regulators failed to trigger significant changes in cell volume in isotonic conditions (Fig. S4c). This indicates that cell volume is not directly governed by cytoskeletal forces. We observed that LatA treatment lowered the cytoplasmic pH after hypotonic shock in 3T3 while having a limited impact on HT1080 (Fig. 3k and Fig. S3i). In addition, 3T3 cells grown on compliant substrates exhibited lower intracellular pH than those grown on glass, implying reduced NHE activity on soft substrates (Fig. S4d). These observations suggest that actomyosin remodeling is required for NHE1 activation during SVI, leading us to postulate that actomyosin indirectly regulates the cell volume by mechanically sensing hypotonic stress and signaling to NHE1, rather than directly exerting forces on the cell cortex and membrane (*22, 34, 42*).

### Mechanosensitive Ca^2+^ signaling regulates cytoskeletal activation of NHE1

To verify our observation that cell volume is regulated by actomyosin mediated mechanosensation, we examined intracellular Ca^2+^ dynamics, which is known to up-regulate myosin activity through Rho kinase and myosin light chain kinase (*53–55*). Using the Ca^2+^ indicator Gcamp6s, we observed that 3T3 cells grown on glass immediately exhibited a greater cytosolic Ca^2+^ increase than HT1080 cells after hypotonic shock (Fig. 4a, b). Soft substrates markedly attenuated such Ca^2+^ increase in 3T3. Hypotonic shock also triggered more high-frequency Ca^2+^ spikes in 3T3 cells on glass compared to those in HT1080 grown on glass (Fig. 4c, d and Fig. S5a, b). Similarly, 3T3 cells on soft substrates exhibited fewer spikes (Fig. 4d and Fig. S5c). Chelating extracellular Ca^2+^ using BAPTA and intracellular Ca^2+^ using BAPTA-AM (Fig. S5d, e), as well as blocking mechanosensitive Ca^2+^ channels using GdCl_3_ or Ruthenium-red (R-red) (Fig. S5f, g), each independently removed SVI and reduced the steady state cell volume in 3T3. We validated a similar effect of Ca^2+^ channel blocking on removing SVI in MCF-10A (Fig. S5j). On the other hand, Ca^2+^ channel blockers had limited impact on the RVD response in HT1080 cells (Fig. S5k,l). These results confirm that Ca^2+^ influx via mechanosensitive ion channels is essential for generating the SVI, and this mechanism is absent in HT1080 cells.

**Figure 4:**
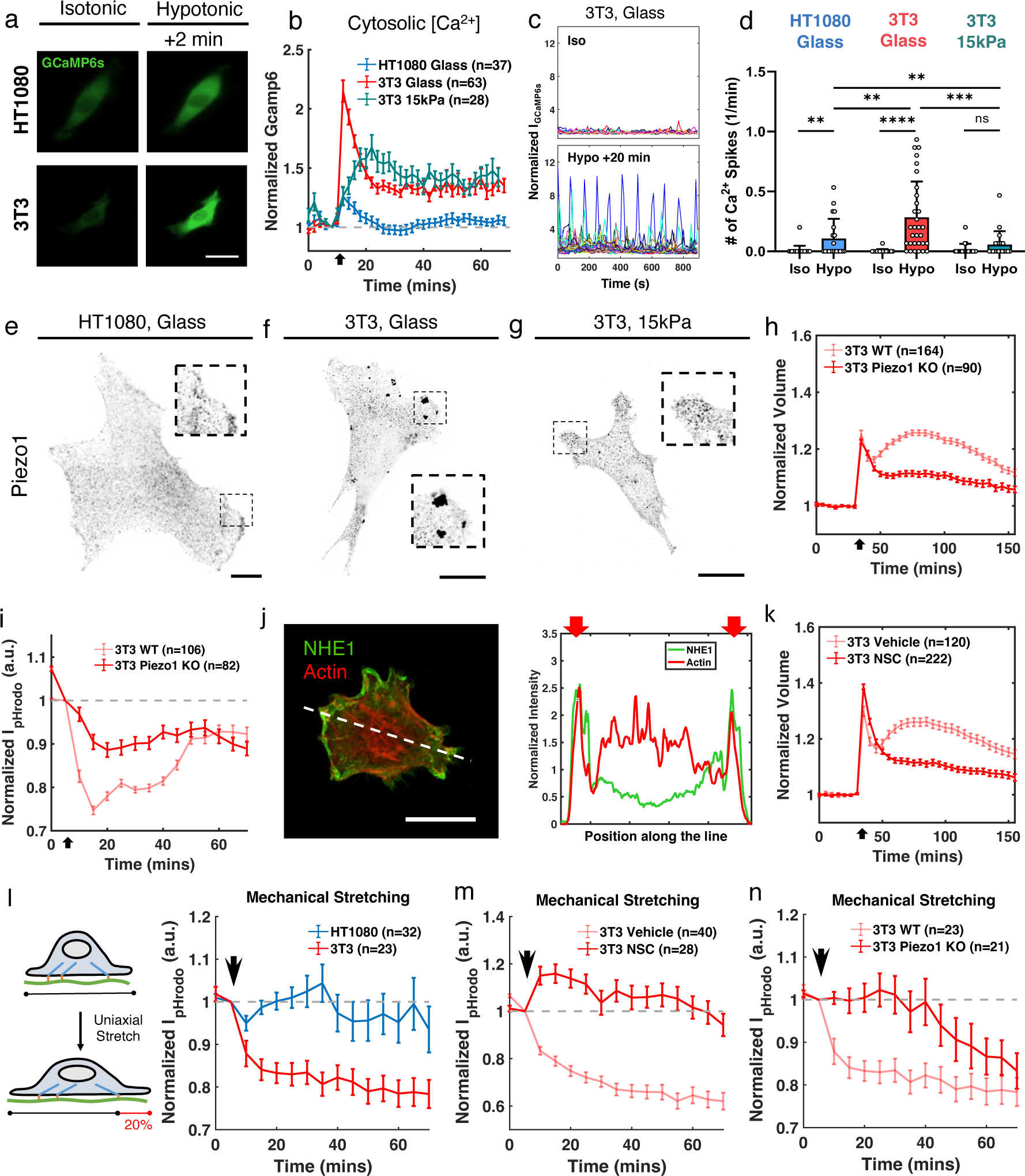
Mechanosensitive Ca^2+^ dynamics regulates cell volume and pH via actin-NHE1 binding partner ezrin. (a) Representative 3T3 and HT1080 GCaMP6s snapshots before and after hypotonic shock. (b) Cytosolic Ca^2+^ concentration of HT1080 grown on glass, 3T3 grown on glass, and 3T3 grown on 15 kPa PDMS substrates measured by GCaMP6s intensity under hypotonic shock (applied at 10 min). Imaged at 2 min frame rate. (c) 3T3 single-cell GCaMP6s dynamics in isotonic media (upper) and 20 min after hypotonic shock (lower) on glass. Imaged at 10 s frame rate. (d) The number of Ca^2+^ spikes in isotonic media versus 20-40 min under hypotonic shock for each cell type and substrate; defined as at least 2x intensity increase over baseline. n = 26, 20, 18, 31, 21, 20, respectively. Mann–Whitney U-tests. (e-g) Immunostaining of Piezo1 at the basal plane for for all cell types and conditions in (d). (h-i) Volume and pHrodo-red AM dynamics in 3T3 WT and Piezo1 KO during hypotonic shock. (j) Co-staining of NHE1 and Actin and their distribution along the line. (k) NSC treated 3T3 volume dynamics under hypotonic shock. (l-n) Normalized pHrodo-red AM intensity of 3T3 versus HT1080 (l), NSC treated 3T3 cells (m), and 3T3 Piezo1 KO (n) under 20% uniaxial mechanical stretch (applied at 5 min). (b, h, i, k-n) Error bars indicate SEM. Mann–Whitney U-tests were used comparing two conditions at each time points after osmotic shock or mechanical stretch, with p values plotted in Supplementary Text 3. (d) Error bars indicate standard deviation. (a, e-g, j) Scale bars, 20 µm.

As one of the most highly expressed Ca^2+^ channels in 3T3 cells (Supplementary Text-1), Piezo1 has been shown to regulate the Ca^2+^ influx triggered by membrane surface expansion (*56–58*). We thus hypothesized that Piezo1 mediates SVI. Immunostaining of Piezo1 showed that 3T3 grown on glass formed dense Piezo1 clusters on the basal plane (Fig. 4e). In contrast, HT1080 grown on glass and 3T3 grown on soft substrates displayed a dispersed Piezo1 localization (Fig. 4f, g), which is unfavorable for its focal adhesion-dependent activation (*59*). CRISPR knockout of Piezo1 in 3T3 (3T3 Piezo1 KO) abolished SVI (Fig. 4h) and the intracellular pH decrease (Fig. 4i) when subjected to hypotonic shock, suggesting that this molecular intervention suppresses NHE1 activity. To validate the role of Piezo1 in volume regulation, we activated Piezo1 using its agonist Yoda1 when applying hypotonic shock. Both HT1080 and MDA-MB-231 cells displayed faster initial RVD (within 3 min) and SVI (Fig. S5h, m). Overall, our results indicate that Ca^2+^ influx through the mechanosensitive ion channel Piezo1 is indispensable for SVI during volume regulation triggered by hypotonic stress.

We also asked how actomyosin communicates with NHE1. We hypothesized that the interaction is facilitated by ezrin, which crosslinks F-actin to NHE1 at the cell membrane (*60, 61*). The co-localization of actin and NHE1 was confirmed by the IF co-staining (Fig. 4j). Next, we examined whether such binding is required for the NHE1 activation during SVI. Blocking ezrin-actin binding via NSC668394 (NSC) effectively removed SVI (Fig. 4k). Interestingly, NSC treatment had limited effects on the initial RVD of 3T3, and its effect in HT1080 was also minimal (Fig. S6a). The distinct effect of NSC compared to EIPA treatment was in agreement with our conclusion that SVI is triggered by additional NHE1 activation via actomyosin. This is not required for the initial RVD. This conclusion was further corroborated by monitoring the intracellular pH in 3T3, where NSC treatment lowered the pH during SVI without affecting the pH increase during the initial RVD (Fig. S6b). Taken together, our results revealed a mechanosensitive regulatory pathway of cell volume: hypotonic stress triggers Ca^2+^ influx through mechanosensitive ion channel Piezo1, inducing actomyosin remodeling that further activates NHE1 through the actin-NHE1 synergistic binding partner ezrin, leading to SVI.

Similar to hypotonic shock, expanding the cell surface by mechanical stretch has been reported to trigger Ca^2+^ influx through Piezo1 (*56*), and we postulated it can also activate NHE1 through the same mechanosensitive pathway. Indeed, applying a one-time, 20% uniaxial stretch to 3T3 led to an immediate pH increase (Fig. 4l). On the contrary, mechanical stretch did not alter the pH in HT1080. Consistent with the hypotonic shock experiments, knocking out Piezo1 or treating 3T3 with NSC or LatA abolished or reduced the immediate pH change after mechanical stretch (Fig. 4m, n, and Fig. S6c). These results support the idea that NHE1 can be mechanically activated by Ca^2+^ and the actin cytoskeleton.

Calmodulin (CaM) is a ubiquitous Ca^2+^ receptor that has been shown to bind to NHE1 (*36, 37, 62*). To examine whether Ca^2+^-CaM regulates NHE1 activity, we treated cells with a Ca^2+^-CaM antagonist, W7. Addition of W7 halted F-actin remodeling during hypotonic shock (Fig. S6g). It also markedly accelerated the initial RVD in both 3T3 and HT1080 (Fig. S6d, e). Importantly, W7 treatment, similar to EIPA treatment, abolished SVI and enhanced initial RVD in 3T3. Addition of W7 also significantly lowered the pH in 3T3 following hypotonic shock (Fig. S6f), indicating that Ca^2+^-CaM is also required for NHE1 activation.

### Cytoskeletal activation of NHE1 deforms nucleus and modifies transcriptomic profile

Recent studies have revealed the important role of the nucleus in sensing mechanical stimuli (*63–68*). To examine whether the cell nucleus responds to hypotonic stress, we stained cells with Hoechst 33342 and monitored nucleus size change during hypotonic shock. HT1080 exhibited an immediate *∼* 10% increase in nuclear area (Fig. 5a, b), consistent with previous reports (*69*). In contrast, 3T3 cells initially exhibited a *∼* 20% reduction in nuclear area, followed by a gradual increase until reaching a similar *∼* 10% increase as HT1080. We noticed that the onset and the timescale of nucleus area recovery coincided with SVI in 3T3, and thus we hypothesized that nucleus deformation is associated with Ca^2+^-cytoskeleton mediated NHE1 activation. Indeed, EIPA treatment hindered the nucleus area recovery in 3T3 (Fig. 5c). Similarly, treating 3T3 with LatA or NSC, or knocking out Piezo1 abolished the nucleus area change during hypotonic shock (Fig. S7a-c).

**Figure 5:**
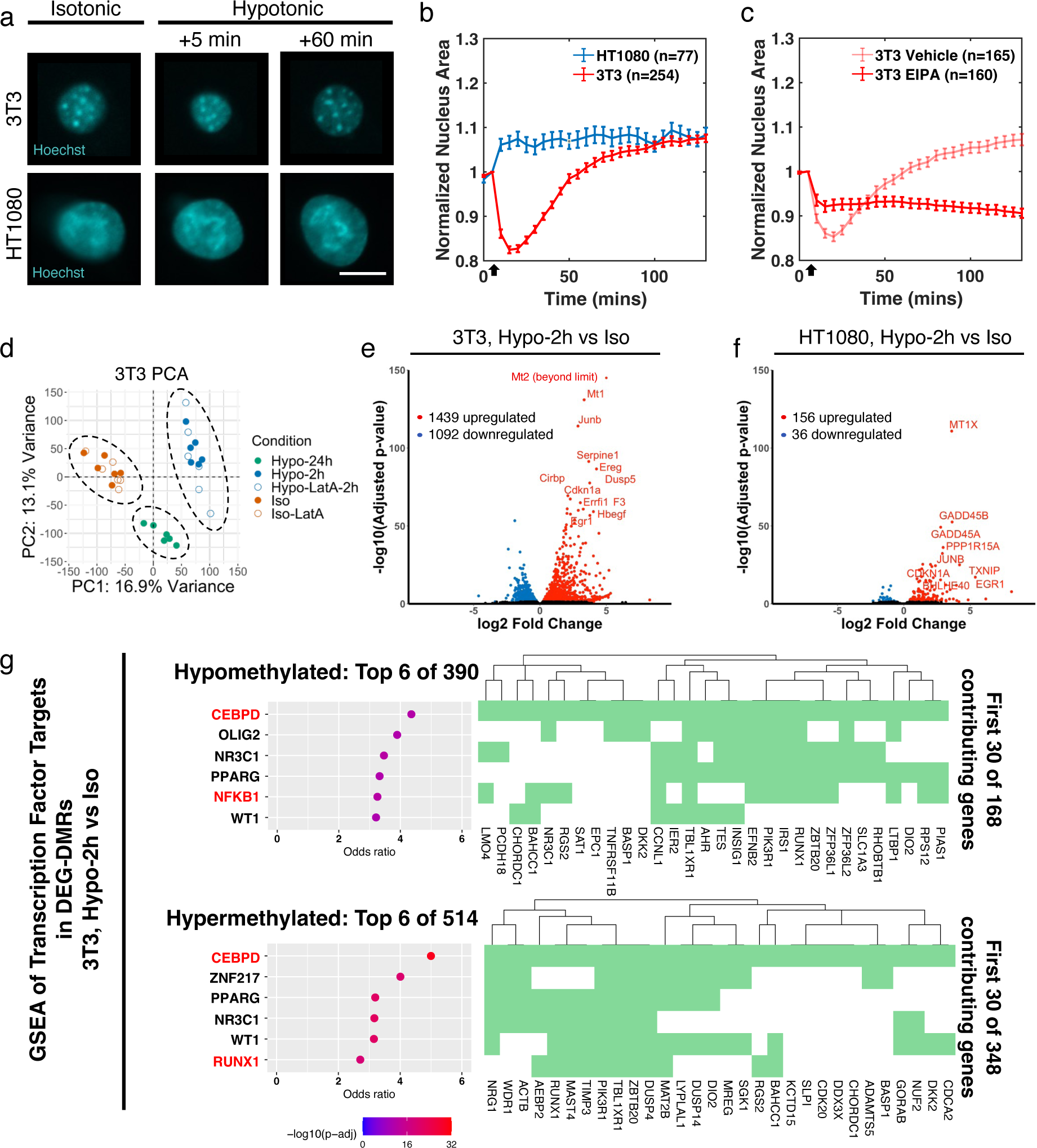
Hypotonic shock deforms cell nucleus and generate large scale transcriptional changes. (a) Representative nuclei images of 3T3 and HT1080 stained with Hoechst 33342 from live single cell tracking. (b) Normalized nucleus area of 3T3 and HT1080 with hypotonic shock applied at 5 min. (c) Normalized nucleus area of EIPA treated 3T3 and vehicle control. Hypotonic shock was applied at 5 min. (d) Principle component analysis (PCA) for 3T3 before and after hypotonic shock and with or without LatA treatment. N = 3 biological repeats. (e, f) Volcano plots showing DEGs at 2-hour hypotonic shock versus in isotonic media for 3T3 (e) and HT1080 (f). (e) Top 6 GSEA terms using ChEA transcription factor database for genes with hypermethylated (top) and hypomethylated (bottom) DEG-DMRs, and top 30 contributing genes (right), at 2-hour hypotonic shock versus in isotonic media for 3T3. (b,c) Error bars indicate SEM. Mann–Whitney U-tests were used comparing two conditions at each time points after osmotic shock, and the p values were plotted in Supplementary Text 3. (a) Scale bars, 10 µm.

The substantial nucleus deformation observed under hypotonic conditions motivated us to further examine the transcriptomic profiles using RNA sequencing (RNA-Seq). Principal component analysis (PCA) clustered all conditions into 3 transcriptomic states: isotonic, 2 h hypotonic shock, and 24 h hypotonic shock, for both 3T3 and HT1080 cell lines regardless of LatA treatment (Fig. 5d and Fig. S7d). Differentially expressed genes (DEGs, located with DESeq2 and defined as adjusted p*<*0.05) between 2 h hypotonic shock and isotonic conditions suggested that 3T3 cells initiated a pronounced, short-timescale transcriptomic response to hypotonic shock that was not observed in HT1080 cells (Fig. 5e, f), in line with their distinct nucleus shape dynamics. Ingenuity Pathway Analysis (IPA, 70) identified the tumor necrosis factor (TNF) signaling pathway as the primary upstream pathway responding to hypotonic shock in both 3T3 and HT1080, but the downstream pathways were distinct in each cell line (Fig. S7g, h). The number of DEGs increased between 24 h after hypotonic shock and isotonic conditions for both cell lines (Fig. S7e, f), indicating hypotonic shock triggered a long-lasting transcriptomic response.

To further explore how cells regulate their transcriptomic state during hypotonic shock, we examined DNA methylation patterns of 3T3 cells via whole-genome bisulfite sequencing (WGBS). Similar to RNA expression, PCA of the WGBS data showed clear segregation into isotonic and hypotonic groups regardless of LatA treatment (Fig. S8a). Differentially methylated regions (DMRs) were located using DMRseq and filtered to identify DMRs over genes with previously identified DEGs in RNA-Seq (termed as DEG-DMRs, Fig. S8b). This analysis revealed 1004 DEG-DMRs between isotonic conditions and 2 h hypotonic shock. Gene set enrichment analysis (GSEA) using the ChEA transcription factor database found significant enrichment of genes with DEG-DMRs for targets of over 900 transcription factors (Fig. 5g). Many top targets are known to be involved in hypoxia including NFKB1, CEBPD, ZNF217, RUNX2, and WT1 (*71–75*), and some of these TFs themselves, notably NFKB1, CEBPD, and RUNX1, have significantly increased expression during hypotonic shock (Fig. S8d). Transcriptional regulation of these TFs and their targets, along with differential DNA methylation at the targets, suggests a robust and highly regulated cellular response to hypotonic shock.

### SVI suppresses ERK/MAPK dependent cell proliferation

We next examined how disrupting SVI using LatA impacted the transcriptome and epigenome. We identified 171 DEGs in 3T3 during 2 h hypotonic shock with LatA treatment compared to the untreated cells (Fig. 6a). IPA revealed several MAPK/ERK related upstream pathways as most significantly inhibited by disrupting SVI (Fig. 6b). Interestingly, LatA treatment suppressed the up-regulation of Dusp2, Dusp5, and Dusp6 observed 2h after hypotonic shock in untreated cells (Fig. 6c). These genes encode phosphatases that inactivate ERK1/2 (*76*), suggesting ERK inhibition is associated with SVI. On the other hand, HT1080 transcriptomics were less sensitive to LatA and no ERK dependence was observed (Fig. S8c). Western blot analysis confirmed that 2 h hypotonic shock reduced ERK phosphorylation in 3T3 WT cells, but not in SVI-deficient 3T3 Piezo1 KO and HT1080 cells (Fig. 6d). Overall, these findings underscore a critical connection between SVI and ERK/MAPK activity.

**Figure 6:**
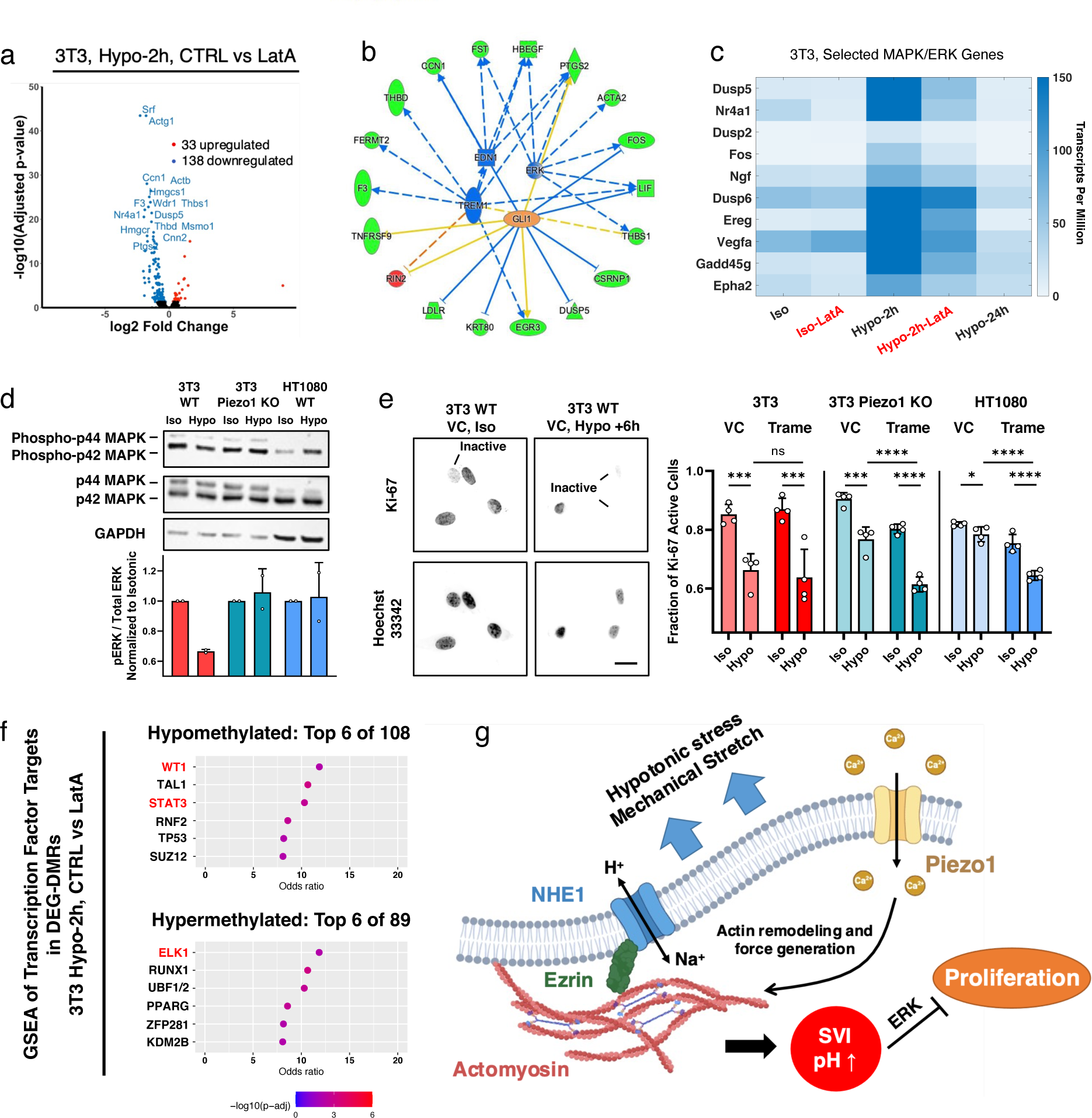
SVI suppress ERK/MAPK dependent cell proliferation. (a) Volcano plots showing DEGs under 2-hour hypotonic shock for between LatA treated 3T3 and vehicle control. (b) Top 4 upstream pathway identified by Ingenuity Pathway Analysis in 3T3 under 2-hour hypotonic shock with and without LatA treatment. Red/green indicates increased/decreased measurement, and orange/blue indicates predicated activation/deactivation. (c) Gene heatmap showing expression of MAPK/ERK pathway genes identify in (b). (d) Western blot for phopho-ERK and total ERK of 3T3 WT, 3T3 Piezo1 KO, and HT1080 cells, with quantification of p-ERK/total ERK ratio. N = 2 biological replicates. (e) Images and quantification of Ki-67 active cells in isotonic conditions and after 6 h of hypotonic shock, with and without Trame treatment in 3T3 WT, 3T3 Piezo1 KO, and HT1080. N = 4 biological replicates, and n *>* 200 cells were examined in each repeat. (f) Top 6 GSEA terms using ChEA transcription factor database for genes with hypermethylated (top) and hypomethylated (bottom) DEG-DMRs at 2-hour hypotonic shock with and without LatA treatment in 3T3. (g) Schematic of the proposed cytoskeletal activation of NHE1 pathway. (d,e) Error bars indicate standard deviation. Mann–Whitney U-tests. (e) Scale bars, 20 µm.

Given that ERK/MAPK signaling pathway is a critical controller of cell proliferation (*77*), we further examined how SVI affected cell proliferation. GSEA identified significant enrichment of targets of ERK related transcription factors in genes with DEG-DMRs between cells treated and untreated with LatA after 2 h hypotonic shock (Fig. 6f). Among them, WT1 has recently been identified to up-regulates ERK/MAPK activity and cell proliferation (*78*), while ELK1 and STAT3 are direct downstream targets of ERK/MAPK that have been shown to control cell growth (*79, 80*). We then used IF to examine the proliferation marker Ki-67 (*81*), and quantify cell proliferation by calculating the fraction of Ki-67 active cells. The fraction of Ki-67 active 3T3 WT cells significantly decreased after 6 h of hypotonic shock. The reduced proliferative potential cannot be further suppressed by ERK inhibition using Trametinib (Trame, Fig. 6e). On the contrary, 3T3 Piezo1 KO and HT1080 cells maintained higher proliferation activities in hypotonic media, which were abolished by ERK inhibition. Taken together, our results revealed that SVI was a suppressor of ERK/MAPK mediated cell proliferation in response to hypotonic stress.

## Discussion

Recent studies have revealed the importance of physical variables such as adhesions and forces in cell size regulation. However, except for a WNK-activated Cl*^−^* secretion mechanism identified during hypertonic volume decrease (*82*), a detailed mechanism of how cells regulate volume during hypotonic expansion has yet to be revealed. In this study, we showed that the initial phase of rapid RVD during hypotonic stress can be explained by rapid actions of ion channels/pumps. However, cells can further regulate their volume by mechanically activating NHE1 through a signaling cascade involving Ca^2+^, actomyosin and ezrin (Fig. 6f), generating SVI and increasing intracellular pH. SVI is largely found in normal-like cell types but is absent in HT1080 and MDA-MB-231 cancer cells, suggesting that differential mechanisms of volume and pH regulation may be a defining feature of transformed and malignant cells. Moreover, the absence of SVI-induced ERK inhibition might facilitate cell proliferation in response to external cues, especially given that ERK-mediated cell proliferation serves as a cancer marker (*77, 83*). While links between mechanosensation, osmoregulation, and cancer have been previously established (*84–86*), the underlying mechanisms remain largely unexplored. Our findings related to SVI and cytoskeletal activation of NHE1 offer a potential bridge between mechanosensation and osmoregulation that could explain the reduced sensitivity of cancer cells under environmental stimuli.

Despite their low concentrations in cytoplasm, our findings highlight the critical role of Ca^2+^ in regulating cell volume and pH through both direct binding and activation of NHE1 via Calmodulin and induction of actomyosin remodel that indirectly activates NHE1 via ezrin. Interestingly, disrupting the actin network or blocking actin-ezrin binding had limited impact on isotonic cell volume (Fig. S4c), suggesting that the cytoskeletal activation of NHE1 observed during SVI and mechanical stretch is a regulatory response to environmental changes, rather than a continuous process during stationary cell culture. Since cells in physiology likely experience varying osmolarities, pressures, and mechanical forces, the mechanosensory system uncovered here is a critical link that connects environmental mechanical cues with cell growth and proliferation.

The magnitude and the timescale of the observed SVI and the corresponding shape change (rounding up) in 3T3 is surprisingly similar to mitotic cell swelling, a phenomenon that has been shown to be regulated by actomyosin and NHE (*10–12*). This correspondence suggests that cells potentially regulate their mitotic entry through Ca^2+^ mediated mechanosensation to ion channels/transporters. Nucleus deformation associated with NHE mediated swelling maybe important for successful mitosis. This hypothesis should be examined in future work, although we note that in HT1080 cells that lack SVI, mitotic swelling is still present. In addition, a series of recent works show that water flux driven by ion transport is responsible for cell migration in 1D confinement and 2D environments, and NHE is one of the central players activated by actomyosin dynamics (*16, 18, 19, 23*). These results establish that the cell volume dynamics is associated with cell migration, and the mechanosensitive activation of NHE1 is another way for cells to generate protrusions and motility. Actin and myosin serve as sensory and control elements during this type of protrusion, but forces are generated by NHE1 mediated water influx to propel the cell. Moreover, the distinct responses to mechanical cues and pharmacological treatments suggest that there are potentially multiple pathways. For example, recent works suggest a potential crosstalk between microtubules and the mTorc pathway that regulates cell volume response mediated by osmotic pressure and curvature of the substrates (*21, 27*). A comprehensive understanding of short timescale cell volume regulation requires a tool that can accurately measure intracellular osmolarity, which is still lacking. We identified the importance of stretch-activated channel Piezo1 in cell volume regulation, but the roles of other mechanosensitive Ca^2+^ channels or store-operated channels need to be explored in the future.

## Materials and Methods

### Cell Culture, osmotic shock media preparation, and pharmacological inhibitors

NIH 3T3, HT1080, and WI-38 cells were a gift from Denis Wirtz (Johns Hopkins University, Baltimore, MD). HEK 293A cells were a gift from Kun-Liang Guan (University of California, San Diego, San Diego, CA). RPE-1 cells were a gift from Rong Li (Johns Hopkins University, Baltimore, MD). MCF-10A, MDA-MB-231, HFF-1, RPE-1 cells were purchased from American Type Culture Collection. 3T3, HT1080, MDA-MB-231, RPE-1, HEK 293A, and HFF-1 were cultured in Dulbecco’s modified Eagle’s media (DMEM; Corning) supplemented with 10% fetal bovine serum (FBS; Sigma), and 1% antibiotics solution contains 10,000 units/mL penicillin and 10,000 µg/mL streptomycin (Gibco). WI-38 cells were cultured in DMEM (low glucose 1g/L; Gibco), with 15% FBS and 1% PS. MCF-10A were cultured in DMEM/F-12 supplemented with 5% horse serum, 20 ng/mL epidermal growth factor, 0.5 µg/mL hydrocortisone, 100 ng/mL cholera toxin, 10 µg/mL insulin, and 1% antibiotics solution, as previously described (*81*). Cells were passaged using 0.05% trypsin-EDTA (Gibco). All cell cultures and live cell experiments were conducted at 37 *^◦^*C and 5% CO2.

Before hypotonic shock experiment, cells were pre-incubated in isotonic solution overnight. The isotonic solution (312 mOsm) contains 50% Dulbecco’s phosphate-buffered saline without calcium and magnesium (Sigma-Aldrich) and 50% cell culture media. 50% hypotonic media (175 mOsm) was prepared by mixing 50% ultra-pure water with 50% cell culture media. For hypertonic shock experiment, cells were cultured in regular cell culture media, and then switched to hypertonic media (504 mOsm). The hypertonic media is the cell culture media supplemented with 160 mM D-Sorbitol (MilliporeSigma). The osmolality was measured using Advanced Instruments model 3320 osmometer.

In select experiments, cells were treated with the following pharmacological agents (purchased from Tocris unless otherwise stated): vehicle controls using DMSO (0.25%, Invitrogen), Latrunculin A (100 nM), Cytochalasin D (500 nM), (S)-nitro-Blebbistatin (1 µM, Cayman Chemical), W7 hydrochloride (20 µM), NSC668394 (10 µM, MilliporeSigma), Ouabain (250 µM), EIPA (100 µM for 3T3, 50 µM for other cell lines), Bumetanide (40 µM), DCPIB (20 µM, Medchemexpress), 4-Aminopyridine (4-AP, 1 mM), Ruthenium red (20 µM), Gadolinium chloride (GdCl3, 20 µM), BAPTA Tetrapotassium Salt (1mM, Invitrogen), BAPTA-AM (5 µM, Invitrogen), and Trametinib (0.2 µM, Medchemexpress). In osmotic shock experiment, cells were pre-adapted in pharmacological agents for 2h expect for Latrunculin A (1 h), Cytochalasin D (1 h), EIPA (1 h), and Ouabain (4 h).

### Microfluidic device fabrication

A detailed protocol can be found in Ref. 40. In brief, FXm channel masks were designed using AutoCAD and ordered from FineLineImaging. Silicon molds were fabricated using SU8-3010 (Kayaku) photoresist following standard photolithography procedures and manufacturer’s protocol. Two layers of photoresist were spin coated on a silicon wafer (IWS) at 500 rpm for 7 s with acceleration of 100 rpm/s, and at 2,000 rpm for 30 s with acceleration of 300 rpm/s, respectively. After a soft bake of 4 min at 95 *^◦^*C, UV light was used to etch the desired patterns from negative photoresist to yield feature heights that were *∼*12 *µ*m. The length of the channels is 16 mm and the width is 1.2 mm.

A 10:1 ratio of PDMS Sylgard 184 silicone elastomer and curing agent were vigorously stirred, vacuum degassed, poured onto each silicon wafer, and cured in an oven at 80 *^◦^*C for 45 min. Razor blades were then used to cut the devices into the proper dimensions, and inlet and outlet ports were punched using a blunt-tipped 21-gauge needle (McMaster Carr; 76165A679). The devices were cleaned by sonicating in 100% isopropyl alcohol for 10 min, and dried using a compressed air gun. The devices and sterilized 50-mm glass-bottom Petri dishes (FlouroDish Cell Culture Dish; World Precision Instruments) were exposed to oxygen plasma for 1 min for bonding. The bonded devices were then placed in an oven at 80 *^◦^*C for 45 min to further ensure bonding.

### Cell volume tracking during pharmacological inhibition and osmotic shock experiment

Micro-fluidic FXm chambers were incubated with 50 µg/mL of type I rat-tail collagen (Enzo) for 1 h at 37 *^◦^*C, followed by washing with isotonic media. Before experiment, *∼* 1-2 million per mL cells with 0.2 mg/mL Alexa Fluor 488 Dextran (MW 2,000 kDa; ThermoFisher) dissolved in isotonic media were injected into the devices using syringe. Cells were then incubated for 1-2 h to allow them to attach and spread. To apply drug or osmotic shock, hypotonic or hypertonic media were injected into the channel gently using a syringe with the same amount of dextran. For osmotic shock experiment with drug treatment, cells were pre-adapted 1-4 h in the FXm device before applying osmotic shock. For cell nucleus area tracking, cells were stained with Hoechst 33342 (1:10000) for 10 min at 37 *^◦^*C in suspension before seeding into the FXm channel. Cells were imaged using fluorescence microscopes as detailed below. The microscope was also equipped with a CO2 module and TempModule stage top incubator (Pecon) that was set to 37 *^◦^*C and 5% CO2 during the experiment. Biological repeat with sample size N *≥* 3 for all experiments.

### Data analysis and cell volume calculation in Fluorescence Exclusion method

Individual cells were tracked by customized MATLAB code using the following algorithm. Firstly, a rough cell mask was draw from a Gaussian blurred epifluorescence image by a set threshold intensity for each cell. The cell mask was then expanded by 100 pixels (2.27 µm) in each direction to ensure proper cell cropping, and a rectangular box was draw based on the largest lateral dimensions in x and y direction. A cell would be discarded if any overlapping cells were found in the box. After all images at different time points were processed, we ran an automatic cell tracking algorithm based on the position of geometric center of each cell mask. In brief, the algorithm assigned each cell (*C_t_*) to the closest cell in the next time frame (*C_t_*_+Δ_*_t_*), and then cross validated the tracking by assigning *C_t_*_+Δ_*_t_* to its closest cell at frame t. The cell would be discarded if the cross validation failed to find *C_t_* as the closest neighbor or found more than one closest neighbor, which is normally due to cell migration, detachment, or overlapping with other cells. The results from automatic cell tracking algorithm were then examined manually by one of the authors.

Cropped images from single cell tracking were then used for cell volume calculation based on the Fluorescence Exclusion method (FXm). The mean fluorescence intensity of the pixels outside cell masks, defined as the mean background intensity *I_bg_*, reflects the height of the FXm channel. The local intensity within the cell mask, *I_V_* defined the difference between channel height and cell height. Given a known channel height *h*, measured by confocal microscopy, the cell volume *V* can be calculated using the equation 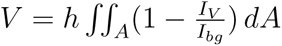

### Epi-fluorescence and confocal microscopy

For epi-fluorescence imaging, a Zeiss Axio Observer inverted, wide-field microscope using a 20× air, 0.8-NA objective or a 63× oil-immersion, 1.2-NA objective equipped with an Axiocam 560 mono charged-coupled device camera was used. In some experiments, a similar microscope equipped with a Hamamatsu Flash4.0 V3 sCMOS camera was used. For confocal imaging, a Zeiss LSM 800 confocal microscope equipped with a 63× oil-immersion, 1.2-NA objective was used. All microscopes were equipped with a CO2 Module S (Zeiss) and TempModule S (Zeiss) stage-top incubator (Pecon) that was set to 37 *^◦^*C with 5% CO2 for live cell imaging. For imaging of immunofluorescence assays, the samples were imaged under room temperature and without CO2. ZEN 2.6 or 3.6 Software (Zeiss) was used as the acquisition software. Customized Matlab (MathWorks) programs or ImageJ were used for image analysis subsequent to data acquisition.

### Measurement of membrane tension using Fluorescence Lifetime Imaging microscopy and Flipper-TR

Cells were seeded into collagen-I coated (50 µg/mL for 1 h at 37 *^◦^*C) glass bottom 24 well plates or 35 mm glass bottom dish coated with 15 kPa PDMS substrates at a density of 5000 cells/cm^2^ in isotonic media overnight. Cells were stained with 1 µM Flipper-TR (Spirochrome) for 30 min and imaged thereafter. Hypotonic shock was then conducted, and cells were imaged 3 min, 30 min, and 2 h after shock. Confocal fluorescence lifetime imaging microscopy was performed using a Zeiss LSM 780 microscope and a PicoQuant system consisting of a PicoHarp 300 time-correlated single-photon counting (TCSPC) module, two-hybrid PMA-04 detectors, and a Sepia II laser control module. A 485 nm laser line was used for excitation with a 600/50 nm band-pass detector unit. Cells were maintained at 37 *^◦^*C with 5% CO2 in a Pecon incubator during imaging. The data analysis was performed as previously described (*47*) using SymPhoTime 64 (PicoQuant) software.

### PDMS substrates fabrication

3 kPa PDMS substrates were prepared by mixing a 1:1 weight ratio of CY52-276A and CY52-276B (Dow) (*24*). To generate 15 kPa and 130 kPa substrates, 2% and 20% Sylgard 527 (Dow, 1:1 base to curing agent ratio) were added to Sylgard 184 (Dow, 10:1 base to curing agent ratio), respectively (*87*). In all cases, the elastomer was vacuum-degassed for approximately 5 min to eliminate bubbles, and then spin-coated onto 35 mm or 50 mm glass bottom dishes at 1,000 rpm for 60 s. The dishes were cured overnight at room temperature, and then at 80 *^◦^*C for 20 min. The devices were subsequently plasma-treated and bonded to the FXm devices for cell volume measurement.

### Cloning, lentivirus preparation, transduction, and transfection

To generate 3T3 cells with stable knockdown of NHE1, the pLKO.1 puro (Addgene; 8453; a gift from Bob Weinberg) backbone was used. For NHE1 depletion, the sequence 5’-GACAAGCTCAACCGGTTTAAT-3’ was subcloned into the pLKO.1 backbone. For generating control cell lines, the non-targeting scramble control sequence 5’-GCACTACCAGAGCTAACTCAGATAGTACT-3’ was subcloned into the pLKO.1 puro backbone.

For lentivirus production, HEK 293T/17 cells were co-transfected with psPAX2, VSVG, and the lentiviral plasmid of interest. 48 h after transfection, the lentivirus was harvested and concentrated using centrifugation. Wild-type 3T3 cells at 60-80% confluency were incubated for 24 h with 100X virus suspension and 8 µg/ml of Polybrene Transfection Reagent (Millipore Sigma). To maintain stable knockdown, cells transduced with the pLKO.1 puro backbone or sg-Bag6 1 with Ef1alpha Puro-P2A-BFP(1)(1) backbone were cultured in medium containing 0.5 µg/ml Puromycin (Gibco).

### Live cell reporters

pEGFP-C1 F-tractin-EGFP (Addgene 58473, a gift from Dyche Mullins) plasmids were used for transient transfections. About 60-80% confluent 3T3 or HT1080 cells were transfected using Lipofectamine 3000 reagent following the manufacturer’s recommendations. pHAGE-RSV-GCaMP6s (Addgene 80146, a gift from Darrell Kotton) plasmids were used for lentiviral cell transduction. The lentivirus production is described above, and cells were selected using flowcytometry.

### CRISPR knockout

Single guide RNA (sgRNA) against PIEZO1 that targets the 5’-AGCATTGAAGCGTAACAGGG-PAM-3’ at Chr.8: 122513787 - 122513809 on GRCm38 was purchased from Invitrogen (A35533). The cells were transfected with the Cas9 enzyme (TrueCut Cas9 Protein v2, Thermo Fisher) and the sgRNA using Lipofectamine™ CRISPRMAX™ Transfection Reagent (Thermo Fisher) according to the manufacturer’s instructions. Transfected cells were then expanded as single-cell clones by limited dilution into 96 well plates. To evaluate the editing efficiency, GeneArt Genomic Cleavage Detection Kit (Life Technologies) was used according to the manufacturer’s instructions. Then, the following primers were used to amplify the region covering the CRISPR binding site: TATCATGGGACCTGGGCATC (forward), CAGGTGTGCACTGAAGGAAC (reverse). The knockout of Piezo1 was confirmed by next generation sequencing (NGS, provided by Genewiz) and western blot (Fig. S5i).

### Immunofluorescence and image analysis

Cells were seeded into collagen-I coated (50 µg/mL for 1 h in 37 °C) glass bottom 24 well plates or 35 mm glass bottom dish coated with 15kPa PDMS substrates at a density of 5000 cells/cm^2^ overnight. For pMLC staining, hypotonic shock was applied and cells were fixed with 4% paraformaldehyde (ThermoFisher Scientific) at different time points as indicated. After washing 3X with PBS, cells were permeabilized in 0.1% Triton X-100 for 15 min. Then, cells were washed 3X in PBS and blocked for 1 h at RT with 5% BSA solution containing 5% normal goat serum (Cell Signaling) and 0.1% Triton X-100. Next, cells were incubated at 4°C with primary antibody overnight. After washing 3X with PBS, samples were incubated for 1 h at RT with secondary antibody. NHE1 staining follows the same protocol as described above but without osmotic shock. Primary antibody: anti-pMLC (Ser19) antibody (Cell Signaling; 3671; 1:100), anti-NHE1 (Invitrogen; PA5-116471; 1:100), anti-Piezo1 (Invitrogen; PA5-77617; 1:500), and anti-Ki-67 (Cell Signaling; 9129; 1:400). Secondary antibody: Alexa Fluor 488 goat anti-rabbit immunoglobulin G (IgG) H+L, (Invitrogen; A11008; 1:200). For actin staining, Alexa Fluor 647 Phalloidin (Invitrogen; A22287; 1:100) were added for 1h at RT. For nucleus staining, Hoechst 33342 (Invitrogen; H3570; 1:10000) were added for 10 min at RT.

Wide-field microscopy using the setup as described above was used to measure the total pMLC and Ki-67contents. A rectangle was cropped for each cell, and the boundary of the cropped area was used for calculating background intensity. The total expressed protein contents of each cell were quantified by summing all intensity after background subtraction. To evaluate Ki-67 expression, cell nucleus masks were drawn based on the Hoechst channel after cropping and background subtraction as aforementioned. The nucleus mask was then applied to Ki-67 staining images to measure the total intensity within each cell nucleus. The spatial distribution of NHE1 and actin was obtained using a confocal microscope as described above. A line was drawn along the cell to obtain the spatial distribution of NHE1 and actin using ImageJ.

### Western Blot

Western blots were performed using protocols previously described in Ref.64, using NuPAGE™ 4-12% Bis-Tris Protein Gels (Thermo Fischer Scientific, NP0336BOX) in Invitrogen Novex NuPage MES SDS Running Buffer (1X, Thermo Fisher Scientific, NP0002). Primary antibody: anti-NHE1 (Santa Cruz Biotechnology; sc-136239; 1:1000), anti-Piezo1 (Invitrogen; PA5-77617; 1:1000), anti-Phospho-p44/42 MAPK (p-ERK1/2; Cell Signaling; 9101; 1:1000), anti-p44/42 MAPK (ERK1/2; Cell Signaling; 4695; 1:1000). GAPDH was used as a loading control (Cell Signaling; 2118; 1:4000). Secondary Antibody: anti-mouse IgG, HRP-linked antibody (Cell Signaling; 7076; 1:1000), and anti-rabbit IgG HRP-linked antibody (Cell Signaling, 7074S; 1:1000).

### Intracellular pH measurement

Intracellular pH was measured using pHrodo Red AM, following the manufacturer’s instructions. Cells were seeded into collagen-I coated (50 µg/mL for 1 h at 37 °C) glass bottom 24 well plates or 35 mm glass bottom dish coated with a 15kPa PDMS substrates at a density of 5000 cells/cm^2^ overnight in isotonic media. Cells were then incubated with 1ml of media containing a dilution of 1 µL of 5 mM pHrodo Red AM in 10 µL of PowerLoad 100X concentrate at 37°C for 30 min. The cells were gently washed once with media and allowed to settle for 15 min before imaging. Cells before and after hypotonic shock were imaged every 5 min using a wide-field epi-fluorescence microscope as previously described. A rectangle larger than the cell size was cropped for each cell, and the signal at the boundary of each cropped area was used to calculate background intensity. pHrodo Red AM signals of individual cells were tracked and measured as the total intensity within the cropped area after background subtraction.

To obtain the absolute pH of different cell types, an intracellular pH calibration buffer Kit (Invitrogen, P35379) was used with pHrodo Red AM, following the manufacturer’s protocol. Briefly, after loading pHrodo Red AM as instructed above, cells were then loaded with the calibration buffer containing 10 µM valinomycin and 10 µM of nigericin at pH values of 4.5, 5.5, 6.7, and 7.5 for at least 5 min. The cells were then imaged and processed through the same background subtraction as described above. Cells were masked, and the mean intensity per pixel was calculated for each cell. A pH calibration curve was then generated by linearly fitting the results from the calibration experiment. Single cell pHrodo intensity was measured, processed, and fitted into the calibration curve to obtain single cell pH results.

### Mechanical stretching

Cells were seeded into elastic PDMS chambers (STB-CH-4W, STREX) at a density of 5000 cells/cm^2^ with cell culture media. The PDMS chambers were pre-coated with 200 µg/mL collagen-I (Enzo) for 4 h at 37 °C. A one-time, 20% uniaxial stretching was applied using a mechanical stretching device (STB-100, STREX). pH measurement and data analysis were conducted as described above. Mechanical stretching may cause cell detachment; therefore, only cells that remained attached and spread both before and after mechanical stretching were analyzed.

### Calcium dynamics imaging and quantification

Cells transiently expressing pGP-CMV-GCaMP6s were seeded into collagen I coated, glass bottom 24 well plate overnight at a density of 5000 cells/cm^2^. For low frequency Ca^2+^ dynamics, cells before and after hypotonic shock were imaged every 2 min using the confocal microscope as described above. Ca^2+^ dynamics were monitored by the mean cell GFP signals per pixel using the image processing method as described above. To measure Ca^2+^ spikes, cells in isotonic media or subjected to hypotonic shock for 20 min were imaged every 10 s using the confocal microscope. Ca^2+^ spikes were identified as instances with greater than 2 times intensity over baseline signals.

### RNA-seq and analysis

Total RNA was extracted and purified using RNeasy Mini Kit (Qiagen) or Quick-DNA/RNA MiniPrep Plus Kit (Zymo Research #D7003) following the manufacturer’s protocol. Strand specific mRNA libraries were generated using the NEBNext Ultra II Directional RNA library prep Kit for Illumina (New England BioLabs #E7760), mRNA was isolated using Poly(A) mRNA magnetic isolation module (New England BioLabs #E7490). Preparation of libraries followed the manufacturer’s protocol (Version 2.2 05/19). Input was 1ug and samples were fragmented for 15 min for RNA insert size of 200 bp. The following PCR cycling conditions were used: 98°C 30s / 8 cycles: 98°C 10s, 65°C 75s / 65°C 5 min. Stranded mRNA libraries were sequenced on an Illumina NovaSeq instrument, SP flowcell using 100bp paired-end dual indexed reads and 1% PhiX control.

Reads were trimmed by 8 base pairs on either end with the Cutadapt package and mapped to the mouse GRCm38 (mm10) or human GRCh38 (hg38) using the Salmon package. Reference genomes were obtained from Ensemble. Batch correction was performed in R with ComBat seq. In R (v.4.2.3), the DESeq2 package was used to determine differentially expressed genes between samples of interest with P values for each gene and comparison. P values were calculated with the DESeq2 package, assuming a negative binomial distribution and correcting for FDR. Differentially expressed genes with an FDR corrected P value *<*= 0.05 were presented in volcano plots and further analyzed. The significantly differentially expressed genes with a log2 fold change *<*= −1 or *>*= 1 were input into Ingenuity Pathway Analysis (Qiagen) to determine pathway enrichment (significance determined by Fischer’s exact method).

### DNA extraction and whole-genome bisulfite sequencing (WGBS)

DNA was extracted using the Quick-DNA/RNA MiniPrep Plus Kit (Zymo Research #D7003) according to the manufacturer’s instructions and genomic DNA samples were quantified by Qubit dsDNA HS assay (ThermoFisher #Q32851). WGBS libraries were prepared using the NEBNext Ultra DNA Library Prep Kit (NEB #E7370L) according to manufacturer’s instructions with modifications. 1% unmethylated Lambda DNA (Promega #D1521) was spiked into genomic DNA to monitor bisulfite conversion efficiency. Genomic DNA (500ng) was fragmented to a target peak of 300-400 bp using a Covaris S2 Focused-ultrasonicator according to the manufacturer’s instructions. Genomic DNA (500 ng) was fragmented to a target peak of 300-400 bp using a Covaris S2 Focused-ultrasonicator according to the manufacturer’s instructions. The fragmented DNA was converted to end-repaired, adenylated DNA using NEBNext Ultra End Repair/dA-Tailing Module (NEB #7442L). Methylated adaptors (NEBNext Multiplex Oligos for Illumina; NEB #E7535L) were ligated to the product from the preceding step using NEBNext Ultra Ligation Module (NEB #7445L). The resulting product was size-selected as described in the manufacturer’s protocol by employing modified SPRIselect (Beckman Coulter #B23318) bead ratios of 0.37X and 0.2X to select for an insert size of 300–400 bp. After size selection, the samples were bisulfite converted and purified using the EZ DNA Methylation-Lightning Kit (Zymo #D5030). Bisulfite-converted libraries were PCR-amplified and uniquely dual-indexed using NEBNext Multiplex Oligos for Illumina (NEB #E6440S) and the Kapa HiFi Uracil+ PCR system (Kapa #KK2801). PCR enrichment was performed with the following cycling parameters: 98°C for 45 sec followed by 8 cycles at 98°C for 15 sec, 65°C for 30 sec, 72°C for 30 sec and a final extension at 72°C for 1 min. The PCR-enriched product was purified via two successive 0.9X SPRIselect bead cleanups. The resulting WGBS libraries were evaluated on a 2100 Bioanalyzer using Agilent High-Sensitivity DNA Kit (Agilent #5067-4626) and quantified via qPCR using the KAPA Library Quantification Kit (KAPA #KK4824). WGBS libraries were sequenced on an Illumina NovaSeq6000 system at a 2 × 150 bp read length with a 5% PhiX control library spike-in.

### Analysis of WGBS data

Adapter sequences were computationally trimmed with Trim Galore, using default parameters for libraries prepared with NEBNext. Bisulfite-aware alignment of the trimmed reads to the mm10 genome was performed using Bismark with default parameters. Samtools was used to merge individual bam files corresponding to fastq file pairs and to name-sort the merged bam file. Methylation bias (mbias) plots were generated using Bismark’s methylation extractor in –mbias only mode. Regions of methylation bias at the 5’ and 3’ ends of reads were determined visually. Bismark was then used to deduplicate merged, name-sorted bam files, and finally, Bismark’s methylation extractor was used to create CpG report files. Successful bisulfite conversion rate was confirmed by alignment of trimmed reads to the lambda genome using Bismark with default parameters.

The Bioconductor package DMRseq was used for DMR-finding. The cytosine report output files from Bismark’s methylation extractor were used as inputs to DMRseq. CpG methylation values were coverage-filtered with a cutoff of 2X for each replicate. Then DMRseq was run to find DMRs; aside from allowing a minimum of 3 CpGs per DMR and using 24 chromosomes per chunk, default parameters were used. Genes overlapping DMRs were identified using Bioconductor package GenomicRanges. DMRs were then filtered to those that overlapped a gene with significant differential expression of RNA. Gene set enrichment analysis was performed using web program Enrichr.

### Statistical analysis

For time series plots, error bars represent the mean and the standard error of mean of at least 3 biological repeats. The number of cells analyzed per condition were noted in the plots or in the captions. For scatter plots, error bar represent the mean and the standard deviation of at least 3 biological repeats. Shapiro–Wilk tests were used for normality testing in cases in which the number of data points was between 3 and 8. D’Agostino–Pearson omnibus normality test was used to determine whether data were normally distributed with *>* 8 data points. For non-Gaussian distributions, nonparametric Mann–Whitney U-tests were used comparing two conditions, and comparisons for more than two groups were performed using Kruskal–Wallis tests followed by Dunn’s multiple-comparison test. For time series plots, Mann–Whitney U-tests were used comparing two conditions at each time points after osmotic shock or mechanical stretch, and the p values were plotted as a function of time in Supplementary Text 3. The statistical analysis was conducted using MATLAB 2021b (Mathworks), or GraphPad Prism 9 or 10 (GraphPad Software). Statistical significance was identified as p*<*0.05. ns for p *>* 0.05, *p*<*0.05, **p*<*0.01, ***p*<*0.001 and ****p*<*0.0001.

## Acknowledgments

We would like to thank Ikbal Choudhury, Shaozheng Sun, Bin Sheng Wong, Wenxuan Du and Rong Li for their helps in experiments or discussions. This work was supported in part by R01 GM134542 (to S.X.S. and K.K.) and NSF 2045715 (to Y.L.). G.S. and A.P.F acknowledge the support from Bloomberg Philanthropies. We also thank John Hopkins University Integrated Imaging Center for microscope support and Rockfish HPC for computational source.

## Author Contributions

Q.N., K.K., and S.X.S conceptualized and designed the study. Q.N., Z.G., and J.F. conducted all live cell experiments and analyzed the data (with Y.Wu). Z.G. and J.F. fabricated all microfluidic devices and PDMS substrates. Y.L. and S.X.S. developed the cell volume model and analyzed the simulation results (with Q.N.). Z.G., Y.Y., A.V. and Y.Wang generated and validated the CRISPR Piezo1-knockout cell line, and K.B. and A.S. generated and validated the shNHE1 cell lines. Q.N., Z.G., A.S., and Y. Y. conducted immunofluorescence and western blots. G.S. and A.P.F. conducted and performed analysis for RNA-Seq and WGBS (with Q.N.). Q.N. and S.X.S. wrote the manuscript, and Z.G., J.F., Y.L., G.S., K.B., A.S., A.P.F., and K.K. edited the manuscript.

## Supplementary Figures

**Figure S1:**
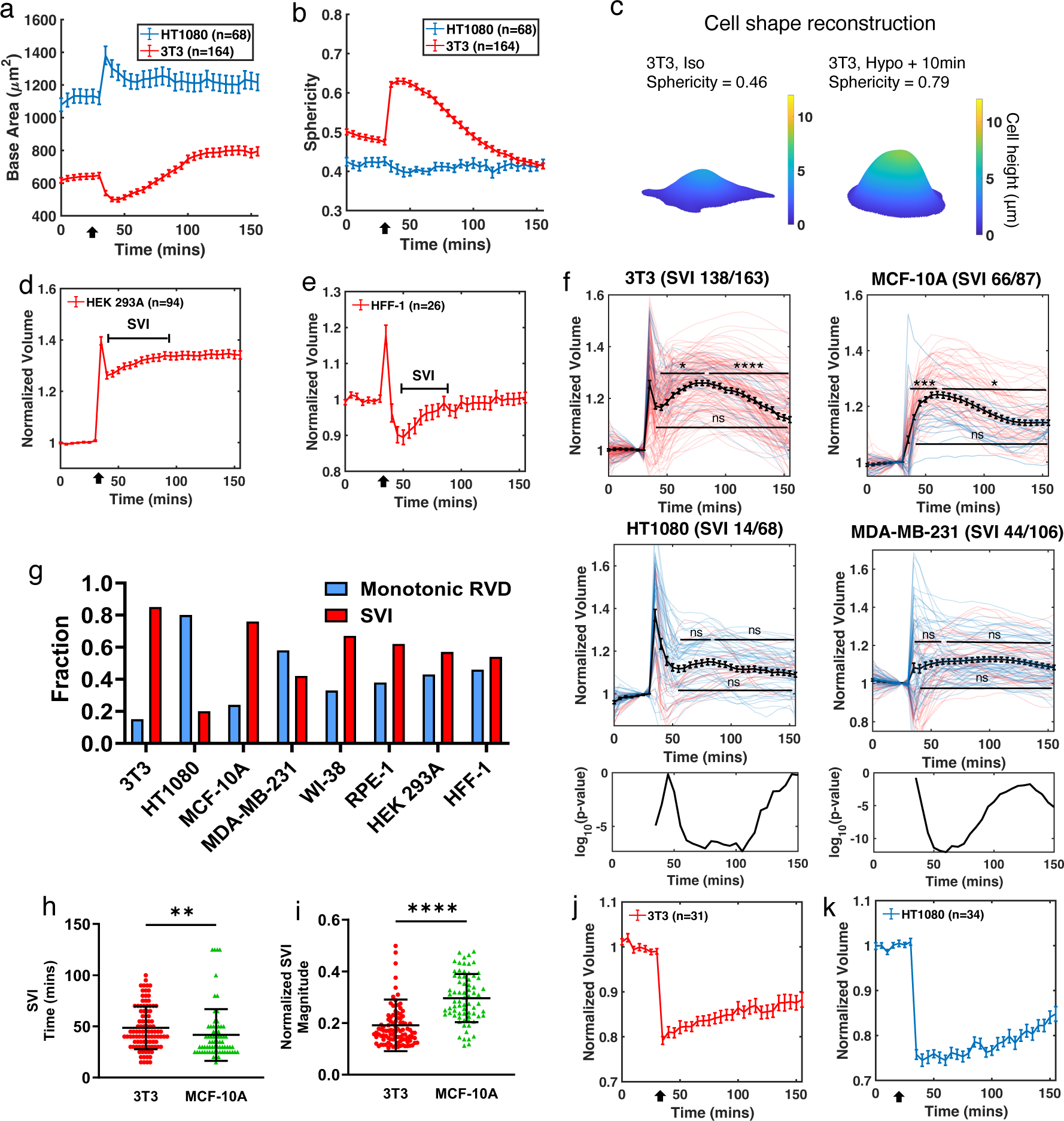
Hypotonic shock induced cell area, shape, and volume changes, SVI variation and statistical significance, and hypertonic shock. (a) Cell base area dynamics during hypotonic shock applied at 30 min. (b) Cell sphericity during hypotonic shock. Sphericity is calculated as 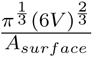. Surface area *A_surface_*is calculated from cell shape reconstruction based on FXm. (c) Representative 3D reconstruction of cell shape from FXm. (d, e) Cell volume tracking of HEK 293A (d) and HFF-1 (e) before and after hypotonic shock. (f) Normalized single cell volume trajectories of 3T3, HT1080, MCF-10A, and MDA-MB-231 before and after hypotonic shock. Cells display at least 10% SVI after initial RVD are considered as a SVI type and are marked in red, and others cells are marked in blue. Mann-Whitney U tests were conducted among the following time points within each cell lines: the first time points after initial RVD (15 min after shock for 3T3, 25 min for HT1080, and 5 min for MCF-10A and MDA-MB-231), the peak of SVI (50 min after shock for 3T3 and HT1080, and 30 min for MCF-10A and MDA-MB-231), and the last time point (2 h after shock). The bottom panels are p-value as a function of time between two pairs of cells: 3T3 versus HT1080 (left) and MCF-10A versus MDA-MB-231 (right) at each time point after shock. ****p*<*0.0001, ***p*<*0.001, **p*<*0.01, and ns p*>*0.05. (g) The fraction of cells showing monotonic RVD and SVI in all cell lines tested. (h, i) The time (h) and magnitude (i) of the SVI in 3T3 and MCF-10A. (j,k) 3T3 (j) and HT1080 (k) volume dynamics with 50% hypotonic shock at 30 min. (a, b, i, j) Error bars indicates standard error of mean (SEM). (f, g) Error bars indicates standard deviation. Mann-Whitney U tests were conducted.

**Figure S2:**
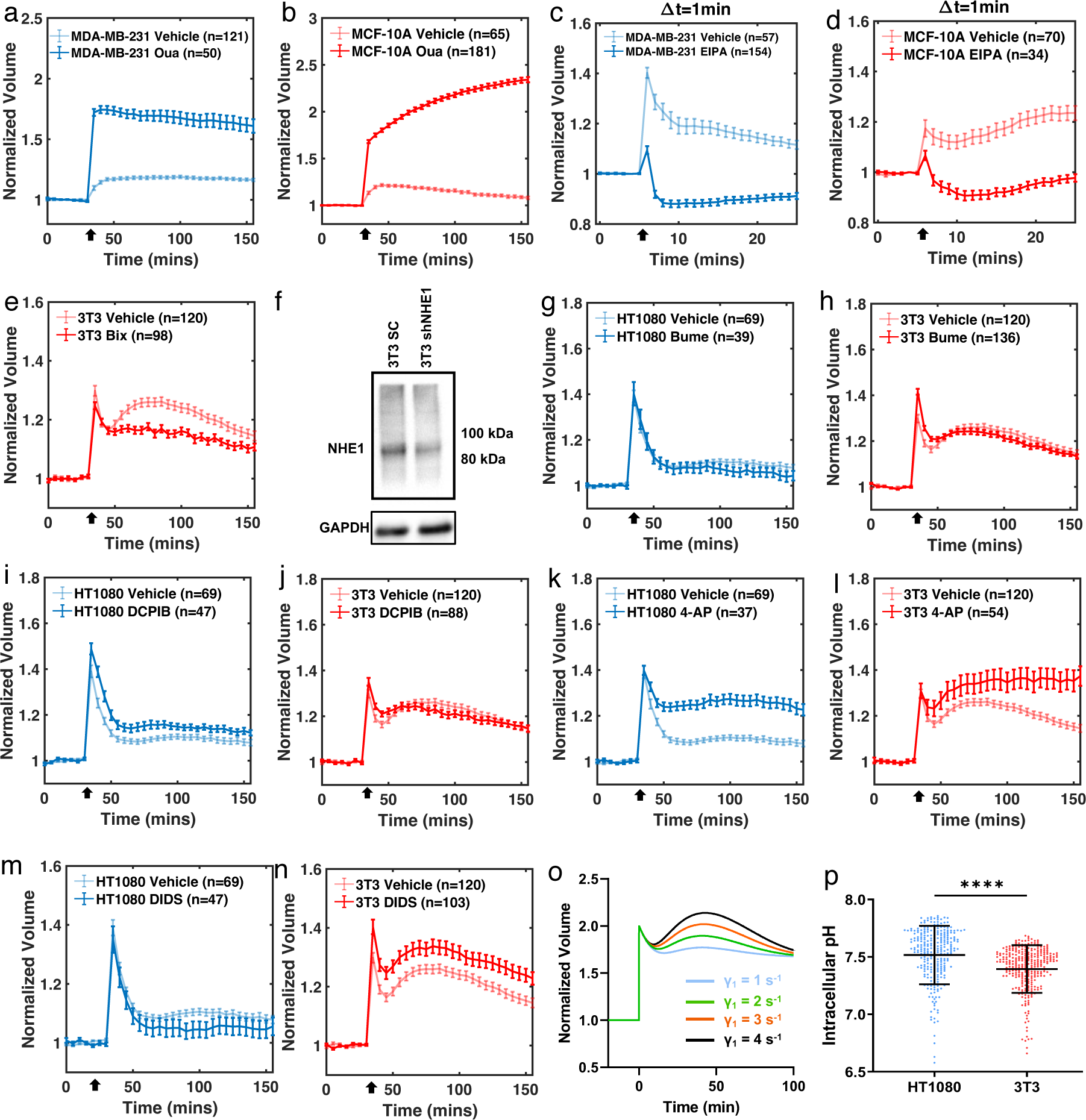
The effects of Na^+^-K^+^ flux in regulating cell volume. (a, b) Oua treated MDA-MB-231 (a) and Mcf-10a (b) volume dynamics under hypotonic shock. (c, d) EIPA treated MDA-MB-231 (c) and Mcf-10a (d) volume dynamics under hypotonic shock (applied at 5 min). Frame rate is 1 min. (e) Bix treated 3T3 volume dynamics under hypotonic shock. (f) Representative western blot from 3T3 transduced with shRNA sequences to NHE1 or a noncoding scramble control (SC) sequence. (g-n) Volume dynamics of 3T3 and HT1080 treated Bume (g for HT1080, h for 3T3), DCPIB (i for HT1080, j for 3T3), 4-AP (k for HT1080, l for 3T3), or DIDS treatment (m for HT1080, n for 3T3). (o) Simulated 3T3 volume dynamics with different NHE activation functions. (p) Intracellular pH of HT1080 and 3T3 at homeostasis. n = 291 and 206, respectively. Error bars indicate standard deviation. Mann-Whitney U tests. (a-e, g-n) Error bars indicate SEM.

**Figure S3:**
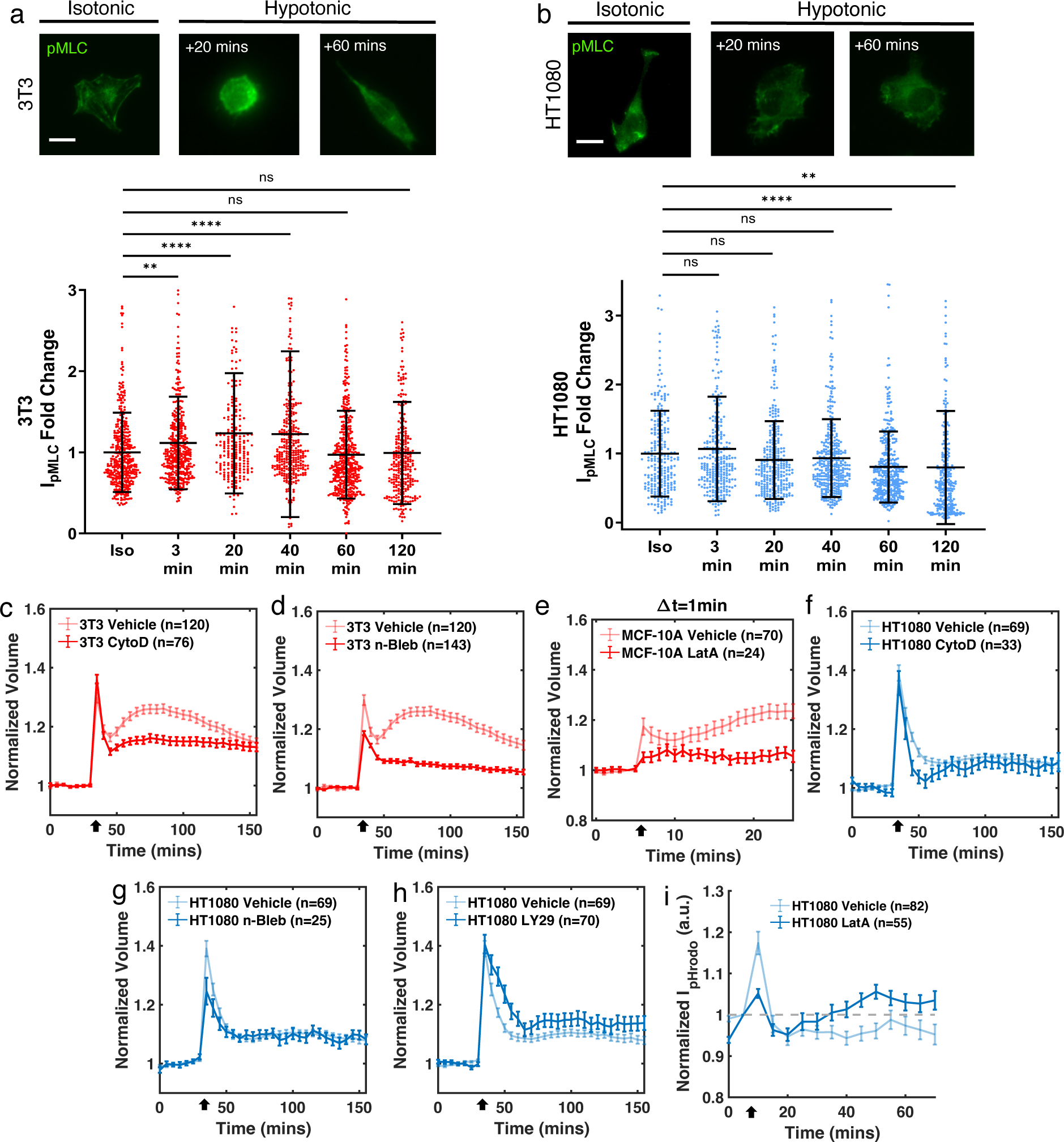
Hypotonic shock induces changes in pohospho-myosin light chain activity and volume dynamics after actomyosin disruption. (a, b) Representative immunofluorescent images of pohospho-myosin light chain and quantifications of 3T3(a) and HT1080 (b). Cells were fixed in isotonic media, and 3, 20, 40, 60 and 120 min after hypotonic shock. For 3T3 (a), n = 405, 401, 216, 330, 441, 287. For HT1080 (b), n = 224, 283, 296, 358, 366, 284. Error bars indicate standard deviation. Kruskal–Wallis tests followed by Dunn’s multiple-comparison test. (c, d) CytoD (c) and n-Bleb (d) treated 3T3 volume dynamics under hypotonic shock. (e) LatA treated MCF-10A volume dynamics under hypotonic shock. Frame rate = 1min. (f-h) CytoD (f), n-Bleb (g), and LY29 (h) trated HT1080 volume dynamics under hypotonic shock. (i) Intensity of pHrodo-red AM for LatA treated HT1080 and vehicle control under hypotonic shock. (c-i) Error bars indicate SEM. (a, b) Scale bars, 20 µm.

**Figure S4:**
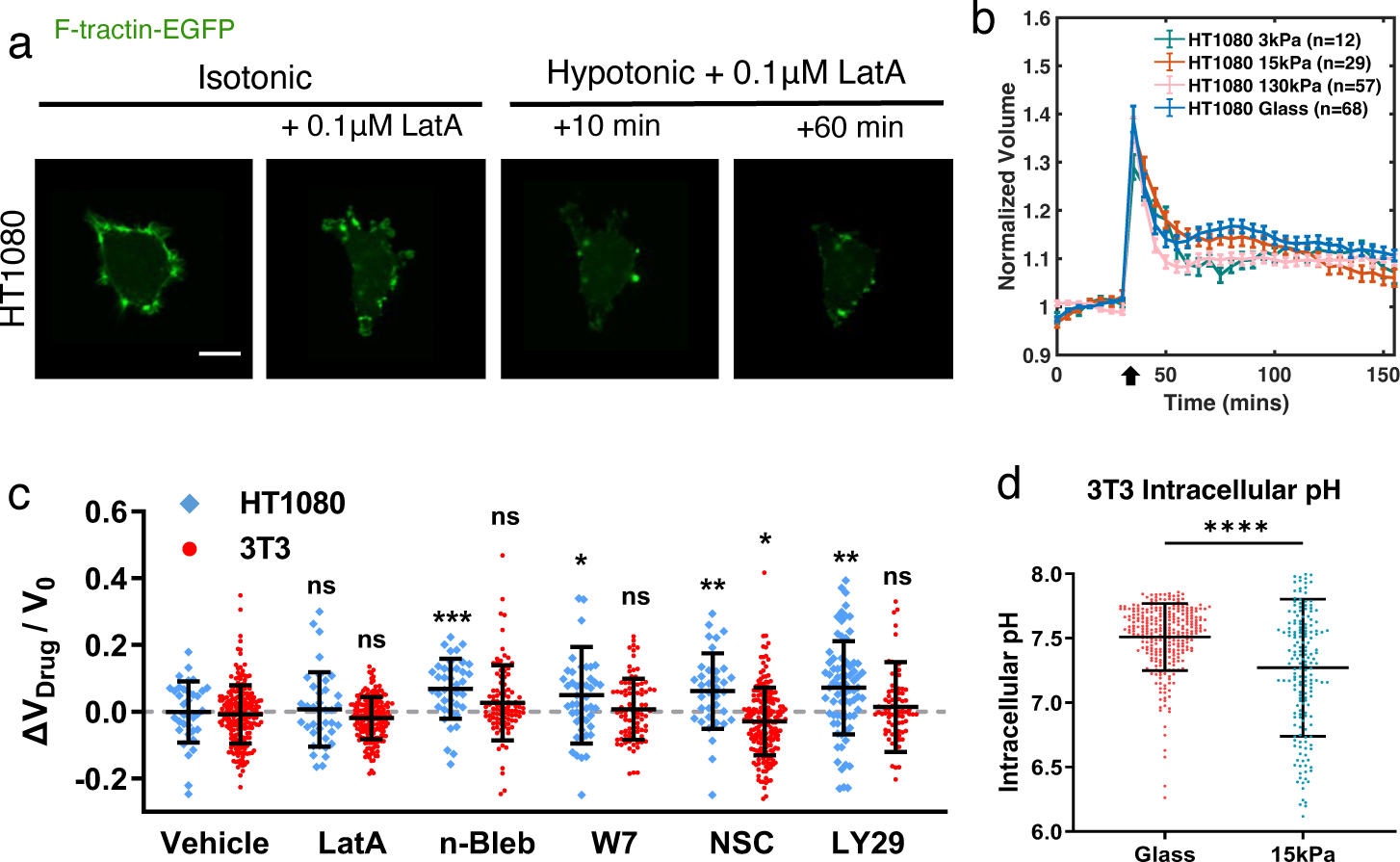
3T3 and HT1080 volume and actin dynamics change under actomyosin disruption or grown on varied substrate stiffness. (a) Confocal images of EGFP-F-tractin transfected HT1080 subjected to LatA treatment before and after hypotonic shock. (b) HT1080 volume dynamics during hypotonic shock on 3, 15, and 130 kPa PDMS substrates and on glass. Error bars indicate SEM. (c) Fractional cell volume changes under vehicle control and drugs for cytoskeleton: Latrunculin A (LatA, dissembling actin network), para-nitro-blebbistatin (n-Bleb, Myosin II inhibitor), W7 (blocking Calmodulin - Ca^2+^ binding), and NSC668394 (NSC, blocking ezrin-actin binding). n_HT1080_ = 36, 190, 40, 46, 36, 77, and n_3T3_ = 213, 38, 102, 89, 202, 84, respectively. Error bars indicate standard deviation. Kruskal–Wallis tests followed by Dunn’s multiple-comparison were conducted between data sets and the vehicle control of the corresponding cell type. (d) Intracellular pH measured by pHrodo-red AM for 3T3 grown on glass and 15kPa PDMS substrates. Error bars indicate standard deviation, and Mann-Whitney U test was conducted. n= 306 and 192, respectively. (a) Scale bar, 20 µm.

**Figure S5:**
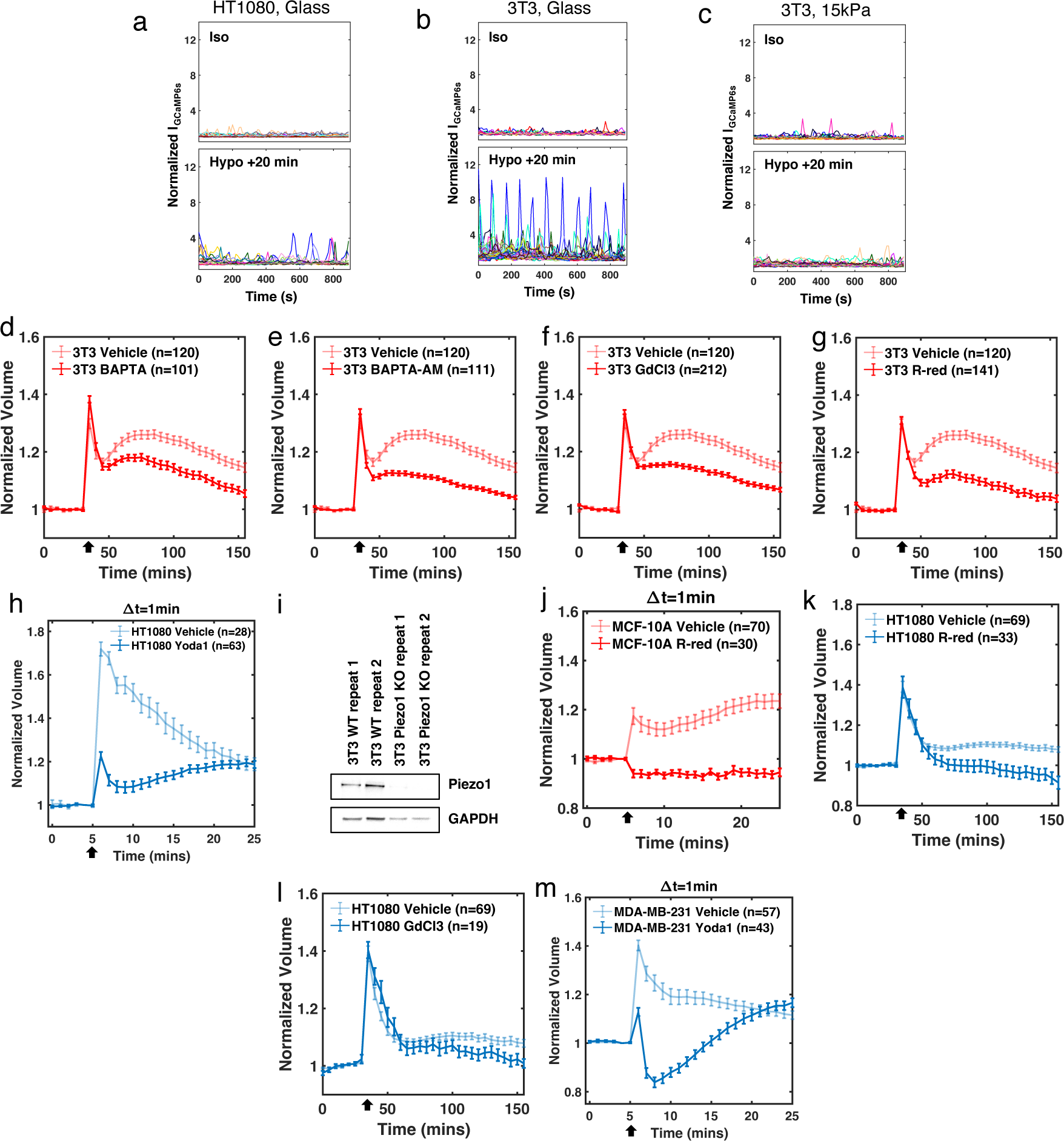
Ca^2+^ signaling affects cell volume regulation. (a-c) Single-cell GCaMP6s dynamics in isotonic media (upper) and 20 min after hypotonic shock (lower) for HT1080 grown on glass (a), 3T3 grown on glass (b), and 3T3 grown on 15 kPa (c). Imaged at 10 s frame rate. (d-g) BAPTA (d), BAPTA-AM (e), GdCl_3_ (f), and R-red (g) treated 3T3 volume dynamics under hypotonic shock. (h) HT1080 volume dynamics with Yoda1 added together with hypotonic solution at 5 min. (i) Representative western blots showing Piezo1 levels in 3T3 WT and Piezo1 CRISPR knockout cells. (j) R-red treated MCF-10A volume dynamics under hypotonic shock applied at 5 min. Frame rate is 1 min. (k-l) R-red (k) and GdCl_3_ (l) treated HT1080 volume dynamics under hypotonic shock. (m) MDA-MB-231 volume dynamics with Yoda1 added together with hypotonic solution at 5 min. (a, d-h, j-m) Error bars indicate SEM.

**Figure S6:**
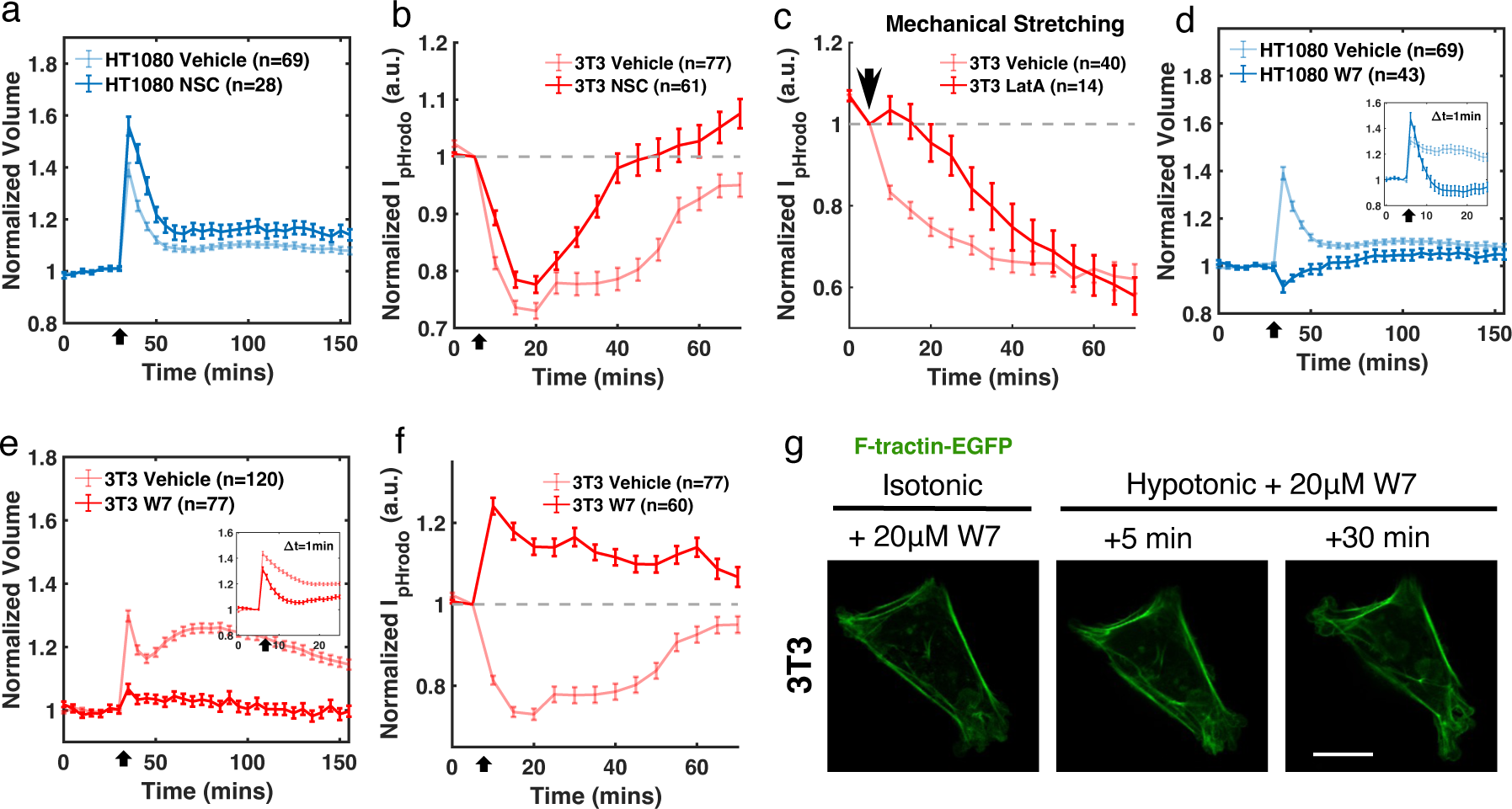
Ca^2+^, calmodulin and ezrin affect cell volume regulation. (a) NSC treated HT1080 volume dynamics under hypotonic shock. (b-c) Intensity of pHrodo-red AM for NSC treated 3T3 under hypotonic shock (b) and 20% uniaxial mechanical stretch (c). (d, e) W7 treated HT1080 (d) and 3T3 (e) volume dynamics under hypotonic shock. Insets have 1 min frame rate (HT1080 Vehicle n = 53, W7 = 20, and 3T3 Vehicle n = 42, W7 n = 46). (f) Intensity of pHrodo-red AM for W7 treated 3T3 under hypotonic shock. (g) 3T3 expressing F-tractin with W7 treatment before and after hypotonic shock. (a-f) Error bars indicate SEM. (g) Scale bar, 20 µm.

**Figure S7:**
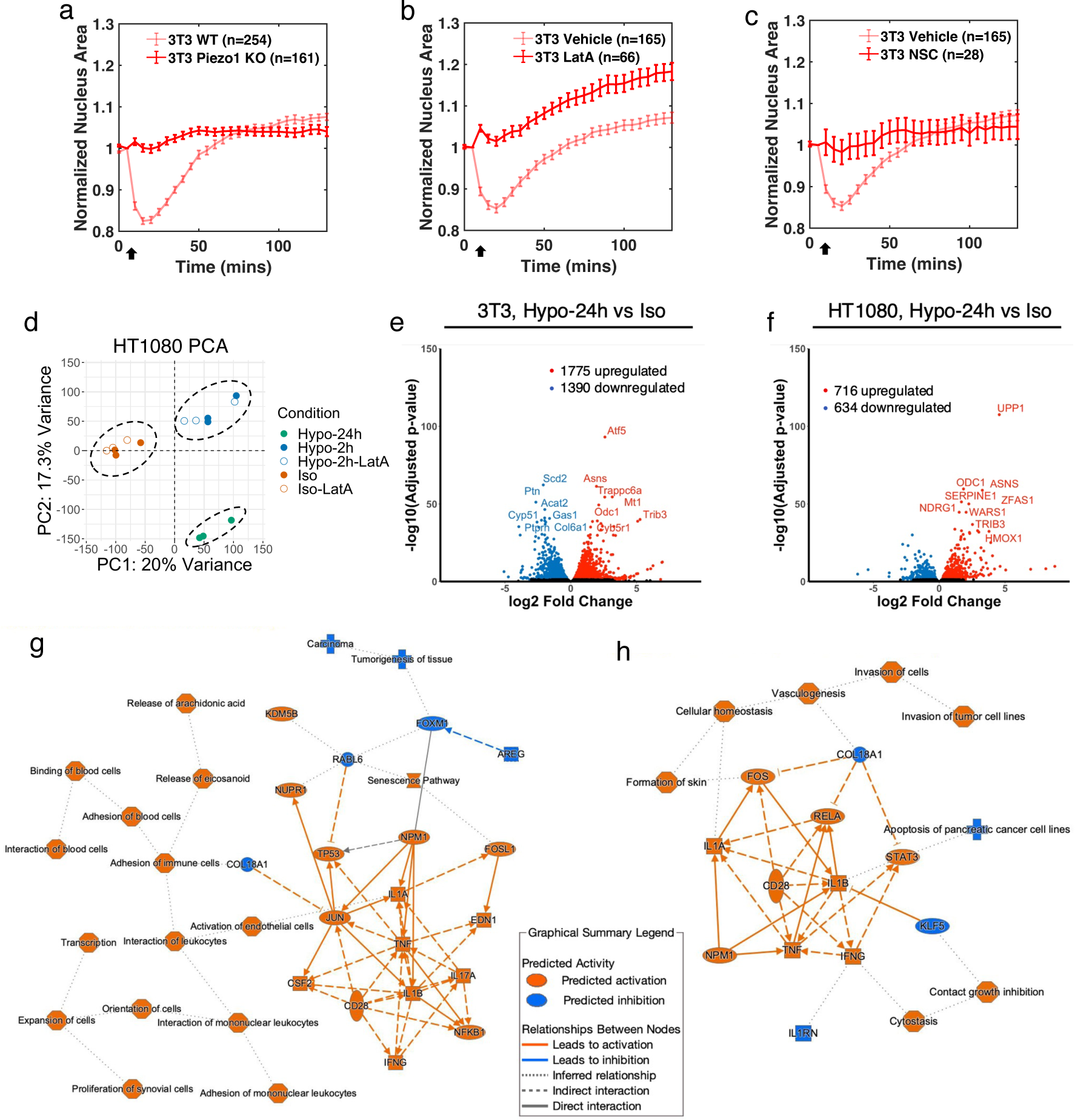
Hypotonic shock induces nucleus deformation and transcriptomic changes. (a-c) Normalized nucleus area of 3T3 WT *versus* Piezo1 KO (a), and 3T3 treated with LatA (b) and NSC (c). Hypotonic shock was applied at 5 min. Error bars indicate SEM. (d) Principle component analysis (PCA) for HT1080 before and after hypotonic shock and with or without LatA treatment. N = 3 biological repeats. (e, f) Volcano plots showing differentially expressed genes (DEGs) at 24-hour hypotonic shock versus in isotonic media for 3T3 (e) and HT1080 (f). (g, h) Summaries of pathways during 2-hour hypotonic shock for 3T3 (g) and HT1080 (h) using Ingenuity Pathway Analysis.

**Figure S8:**
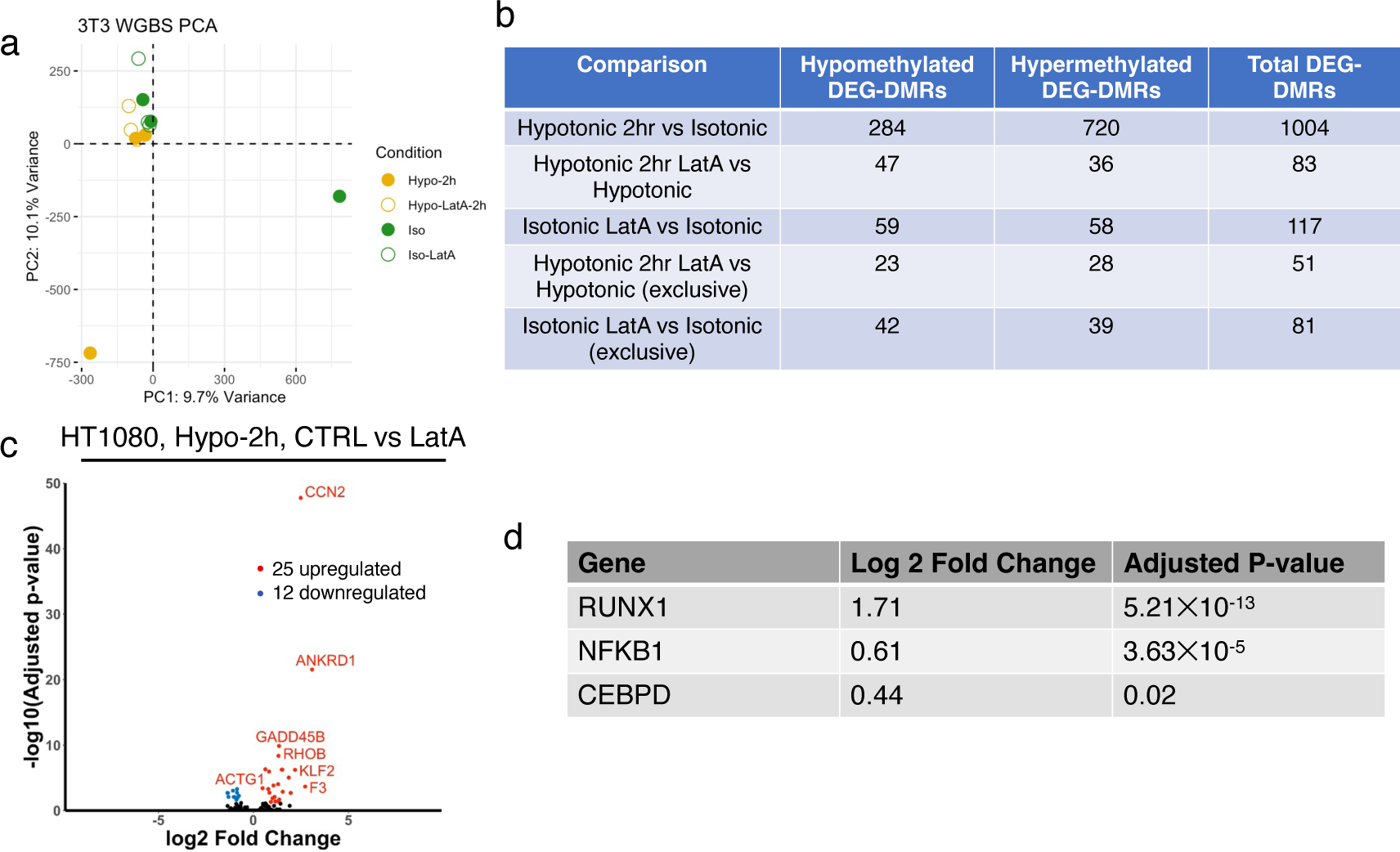
DNA methylation and HT1080 RNAseq. (a) Principle component analysis (PCA) of 3T3 WGBS data before and after 2 h hypotonic shock and with or without LatA treatment. N = 3 biological repeats. (b) Summary of the number of DEG-DMRs comparing all conditions from 3T3 WGBS data. (c) Volcano plots showing DEGs at 2-hour hypotonic shock control versus LatA treatment for HT1080. (d) Differential expression of select transcription factors between Hypotonic 2 h and Isotonic cells.

## Uncropped western blot

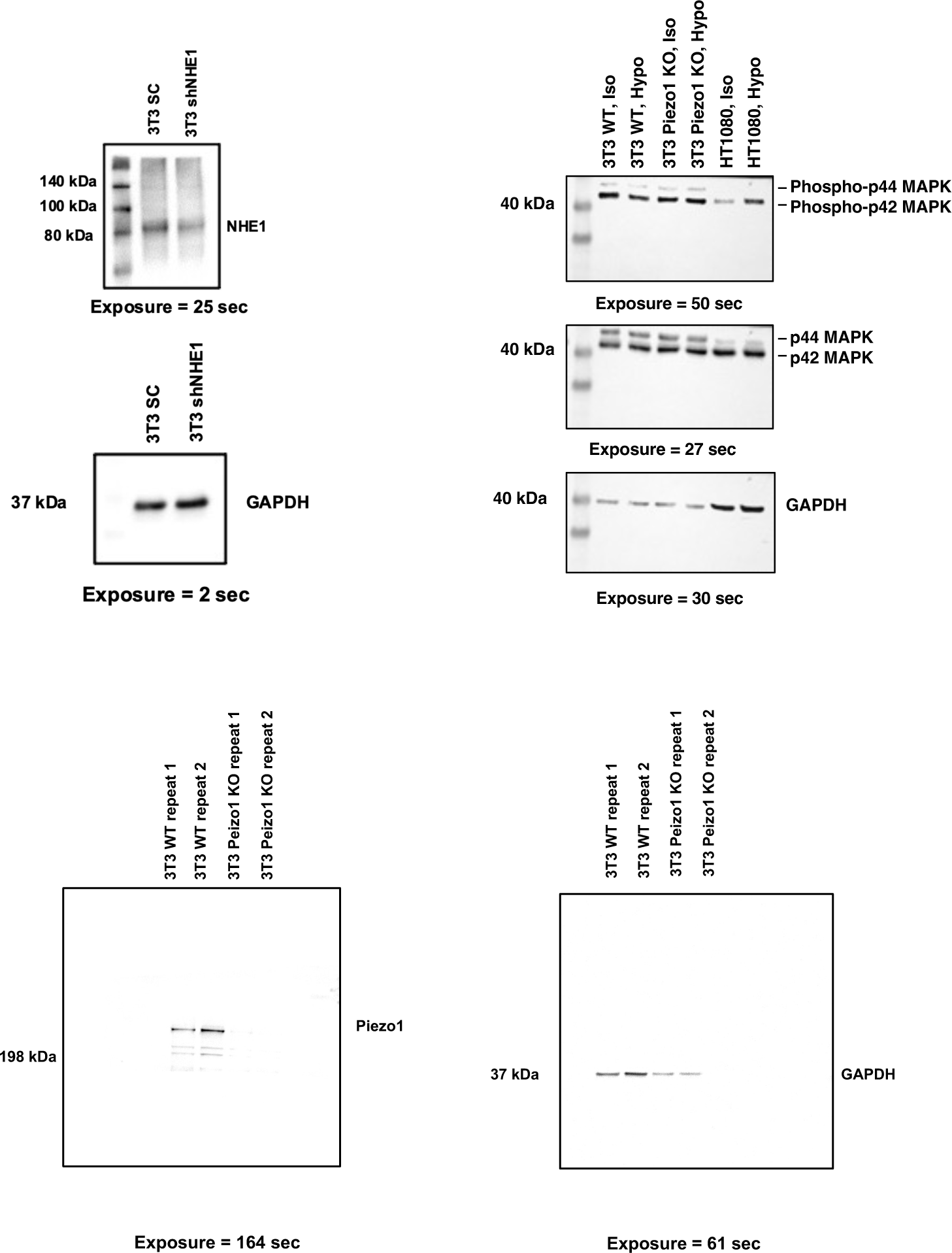

## References

1. Else K. Hoffmann, Ian H. Lambert, and Stine F. Pedersen. Physiology of cell volume regulation in vertebrates. Physiological Reviews, 89(1):193–277, 2009.

2. Miriam B. Ginzberg, Ran Kafri, and Marc Kirschner. On being the right (cell) size. Science, 348(6236), 2015.

3. Hung Ji Tsai, Anjali R. Nelliat, Mohammad Ikbal Choudhury, Andrei Kucharavy, William D. Bradford, Malcolm E. Cook, Jisoo Kim, Devin B. Mair, Sean X. Sun, Michael C. Schatz, and Rong Li. Hypo-osmotic-like stress underlies general cellular defects of aneuploidy. Nature, 570(7759):117–121, 2019.

4. Gabriel E. Neurohr, Rachel L. Terry, Jette Lengefeld, Megan Bonney, Gregory P. Brittingham, Fabien Moretto, Teemu P. Miettinen, Laura Pontano Vaites, Luis M. Soares, Joao A. Paulo, J. Wade Harper, Stephen Buratowski, Scott Manalis, Folkert J. van Werven, Liam J. Holt, and Angelika Amon. Excessive Cell Growth Causes Cytoplasm Dilution And Contributes to Senescence. Cell, 176(5):1083–1097.e18, 2019.

5. Michael C. Lanz, Evgeny Zatulovskiy, Matthew P. Swaffer, Lichao Zhang, Ilayda Ilerten, Shuyuan Zhang, Dong Shin You, Georgi Marinov, Patrick McAlpine, Joshua E. Elias, and Jan M. Skotheim. Increasing cell size remodels the proteome and promotes senescence. Molecular Cell, 82(17):3255–3269.e8, 2022.

6. Clotilde Cadart, Sylvain Monnier, Jacopo Grilli, Pablo J. Sáez, Nishit Srivastava, Rafaele Attia, Emmanuel Terriac, Buzz Baum, Marco Cosentino-Lagomarsino, and Matthieu Piel. Size control in mammalian cells involves modulation of both growth rate and cell cycle duration. Nature Communications, 9(1), 2018.

7. Yufei Wu, Adrian F Pegoraro, David A Weitz, Paul Janmey, and Sean X Sun. The correlation between cell and nucleus size is explained by an eukaryotic cell growth model. PLoS computational biology, 18(2):e1009400, feb 2022.

8. Romain Rollin, Jean-François Joanny, and Pierre Sens. Physical basis of the cell size scaling laws. eLife, 12:1–20, may 2023.

9. Xili Liu, Jiawei Yan, and Marc W Kirschner. Cell size homeostasis is tightly controlled throughout the cell cycle. PLoS biology, 22(1):e3002453, jan 2024.

10. Ewa Zlotek-Zlotkiewicz, Sylvain Monnier, Giovanni Cappello, Mael Le Berre, and Matthieu Piel. Optical volume and mass measurements show that mammalian cells swell during mitosis. Journal of Cell Biology, 211(4):765–774, 2015.

11. Sungmin Son, Joon Ho Kang, Seungeun Oh, Marc W. Kirschner, T. J. Mitchison, and Scott Manalis. Resonant microchannel volume and mass measurements show that suspended cells swell during mitosis. Journal of Cell Biology, 211(4):757–763, 2015.

12. Martin P Stewart, Jonne Helenius, Yusuke Toyoda, Subramanian P Ramanathan, Daniel J Muller, and Anthony A Hyman. Hydrostatic pressure and the actomyosin cortex drive mitotic cell rounding. Nature, 469(7329):226–230, 2011.

13. Hongyuan Jiang and Sean X. Sun. Cellular pressure and volume regulation and implications for cell mechanics. Biophysical Journal, 105(3):609–619, 2013.

14. Jiaxiang Tao and Sean X. Sun. Active Biochemical Regulation of Cell Volume and a Simple Model of Cell Tension Response. Biophysical Journal, 109(8):1541–1550, oct 2015.

15. Oscar M Lancaster, Maël Le Berre, Andrea Dimitracopoulos, Daria Bonazzi, Ewa Zlotek- Zlotkiewicz, Remigio Picone, Thomas Duke, Matthieu Piel, and Buzz Baum. Mitotic rounding alters cell geometry to ensure efficient bipolar spindle formation. Developmental cell, 25(3):270–83, may 2013.

16. Kimberly M. Stroka, Hongyuan Jiang, Shih Hsun Chen, Ziqiu Tong, Denis Wirtz, Sean X. Sun, and Konstantinos Konstantopoulos. Water permeation drives tumor cell migration in confined microenvironments. Cell, 157(3):611–623, 2014.

17. Jiaxiang Tao, Yizeng Li, Dhruv K. Vig, and Sean X. Sun. Cell mechanics: A dialogue. Reports on Progress in Physics, 80(3), 2017.

18. Yizeng Li, Konstantinos Konstantopoulos, Runchen Zhao, Yoichiro Mori, and Sean X. Sun. The importance of water and hydraulic pressure in cell dynamics. Journal of cell science, 133(20), 2020.

19. Kaustav Bera, Alexander Kiepas, Inês Godet, Yizeng Li, Pranav Mehta, Brent Ifemembi, Colin D Paul, Anindya Sen, Selma A Serra, Konstantin Stoletov, Jiaxiang Tao, Gabriel Shatkin, Se Jong Lee, Yuqi Zhang, Adrianna Boen, Panagiotis Mistriotis, Daniele M Gilkes, John D Lewis, Chen-Ming Fan, Andrew P Feinberg, Miguel A Valverde, Sean X Sun, and Konstantinos Konstantopoulos. Extracellular fluid viscosity enhances cell migration and cancer dissemination. Nature, 611(November), 2022.

20. Clotilde Cadart, Larisa Venkova, Pierre Recho, Marco Cosentino Lagomarsino, and Matthieu Piel. The physics of cell-size regulation across timescales. Nature Physics, 15(10):993–1004, 2019.

21. Chloé Roffay, Guillaume Molinard, Kyoohyun Kim, Marta Urbanska, Virginia Andrade, Victoria Barbarasa, Paulina Nowak, Vincent Mercier, José García-Calvo, Stefan Matile, Robbie Loewith, Arnaud Echard, Jochen Guck, Martin Lenz, and Aurélien Roux. Passive coupling of membrane tension and cell volume during active response of cells to osmosis. Proceedings of the National Academy of Sciences of the United States of America, 118(47), 2021.

22. Larisa Venkova, Amit Singh Vishen, Sergio Lembo, Nishit Srivastava, Baptiste Duchamp, Artur Ruppel, Alice Williart, Stéphane Vassilopoulos, Alexandre Deslys, Juan Manuel Garcia Arcos, Alba Diz-Muñoz, Martial Balland, Jean-François Joanny, Damien Cuvelier, Pierre Sens, and Matthieu Piel. A mechano-osmotic feedback couples cell volume to the rate of cell deformation. eLife, 11, apr 2022.

23. Debonil Maity, Kaustav Bera, Yizeng Li, Zhuoxu Ge, Qin Ni, Konstantinos Konstantopoulos, and Sean X. Sun. Extracellular Hydraulic Resistance Enhances Cell Migration. Advanced Science, 9(29):2200927, oct 2022.

24. Nicolas Perez Gonzalez, Jiaxiang Tao, Nash D. Rochman, Dhruv Vig, Evelyn Chiu, Denis Wirtz, and Sean X. Sun. Cell tension and mechanical regulation of cell volume. Molecular Biology of the Cell, 29(21):2591–2600, 2018.

25. Ming Guo, Adrian F. Pegoraro, Angelo Mao, Enhua H. Zhou, Praveen R. Arany, Yulong Han, Dylan T. Burnette, Mikkel H. Jensen, Karen E. Kasza, Jeffrey R. Moore, Frederick C. Mackintosh, Jeffrey J. Fredberg, David J. Mooney, Jennifer Lippincott-Schwartz, and David A. Weitz. Cell volume change through water efflux impacts cell stiffness and stem cell fate. Proceedings of the National Academy of Sciences of the United States of America, 114(41):E8618–E8627, 2017.

26. Kenan Xie, Yuehua Yang, and Hongyuan Jiang. Controlling Cellular Volume via Mechanical and Physical Properties of Substrate. Biophysical Journal, 114(3):675–687, 2018.

27. Caterina Tomba, Valeriy Luchnikov, Luca Barberi, Carles Blanch-Mercader, and Aurélien Roux. Epithelial cells adapt to curvature induction via transient active osmotic swelling. Developmental Cell, 57(10):1257–1270.e5, may 2022.

28. T. H. Hui, Z. L. Zhou, J. Qian, Y. Lin, A. H.W. Ngan, and H. Gao. Volumetric deformation of live cells induced by pressure-activated cross-membrane ion transport. Physical Review Letters, 113(11):1–5, 2014.

29. Jiwon Park, Siyang Jia, Donald Salter, Pierre Bagnaninchi, and Carsten G Hansen. The Hippo pathway drives the cellular response to hydrostatic pressure. The EMBO Journal, pages 1–21, 2022.

30. Stacey Watkins and Harald Sontheimer. Hydrodynamic cellular volume changes enable glioma cell invasion. The Journal of neuroscience : the official journal of the Society for Neuroscience, 31(47):17250–9, nov 2011.

31. D C Tosteson and J F Hoffman. Regulation of cell volume by active cation transport in high and low potassium sheep red cells. The Journal of general physiology, 44:169–94, sep 1960.

32. Horacio F. Cantiello. Role of actin filament organization in cell volume and ion channel regulation. The Journal of experimental zoology, 279(5):425–35, dec 1997.

33. John H. Henson. Relationships between the actin cytoskeleton and cell volume regulation. Microscopy Research and Technique, 47(2):155–162, 1999.

34. Ram M Adar and Samuel A Safran. Active volume regulation in adhered cells. Proceedings of the National Academy of Sciences of the United States of America, 117(11):5604–5609, mar 2020.

35. Ram M Adar, Amit Singh Vishen, Jean-François Joanny, Pierre Sens, and Samuel A Safran. Volume regulation in adhered cells: Roles of surface tension and cell swelling. Biophysical journal, 122(3):506–512, feb 2023.

36. Xiuju Li, Daniel Prins, Marek Michalak, and Larry Fliegel. Calmodulin-dependent binding to the NHE1 cytosolic tail mediates activation of the Na+/H+ exchanger by Ca2+ and endothelin. American Journal of Physiology - Cell Physiology, 305(11):1161–1169, 2013.

37. Lise M Sjøgaard-Frich, Andreas Prestel, Emilie S Pedersen, Marc Severin, Kristian Kølby Kristensen, Johan G Olsen, Birthe B Kragelund, and Stine Falsig Pedersen. Dynamic Na+/H+ exchanger 1 (NHE1) - calmodulin complexes of varying stoichiometry and structure regulate Ca2+-dependent NHE1 activation. eLife, 10, mar 2021.

38. Lijuan He, Jiaxiang Tao, Debonil Maity, Fangwei Si, Yi Wu, Tiffany Wu, Vishnu Prasath, Denis Wirtz, and Sean X. Sun. Role of membrane-tension gated Ca 2+ flux in cell mechanosensation. Journal of Cell Science, 131(4):1–12, 2018.

39. C. Cadart, E. Zlotek-Zlotkiewicz, L. Venkova, O. Thouvenin, V. Racine, M. Le Berre, S. Monnier, and M. Piel. Fluorescence eXclusion Measurement of volume in live cells. Methods in cell biology, 139:103–120, 2017.

40. Nash Rochman, Kai Yao, Nicolas Perez Gonzalez, Denis Wirtz, and Sean Sun. Single Cell Volume Measurement Utilizing the Fluorescence Exclusion Method (FXm). BIO-PROTOCOL, 10(12), 2020.

41. Thomas J. Jentsch. VRACs and other ion channels and transporters in the regulation of cell volume and beyond. Nature Reviews Molecular Cell Biology, 17(5):293–307, may 2016.

42. Yizeng Li, Xiaohan Zhou, and Sean X Sun. Hydrogen, Bicarbonate, and Their Associated Exchangers in Cell Volume Regulation. Frontiers in Cell and Developmental Biology, 9:1–22, jun 2021.

43. Tamas L Nagy, Jack D Strickland, and Orion D Weiner. Neutrophils actively swell to potentiate rapid migration. eLife, (12:RP90551), 2023.

44. Gerdi Christine Kemmer, Sidsel Ammitzbøll Bogh, Michael Urban, Michael G Palmgren, Tom Vosch, Jürgen Schiller, and Thomas Günther Pomorski. Lipid-conjugated fluorescent pH sensors for monitoring pH changes in reconstituted membrane systems. The Analyst, 140(18):6313–20, sep 2015.

45. Arkady S. Pivovarov, Fernando Calahorro, and Robert J. Walker. Na+/K+-pump and neurotransmitter membrane receptors. Invertebrate Neuroscience, 19(1):1, mar 2019.

46. Farshid Guilak, Geoffrey R Erickson, and H Ping Ting-Beall. The effects of osmotic stress on the viscoelastic and physical properties of articular chondrocytes. Biophysical journal, 82(2):720–7, feb 2002.

47. Adai Colom, Emmanuel Derivery, Saeideh Soleimanpour, Caterina Tomba, Marta Dal Molin, Naomi Sakai, Marcos González-Gaitán, Stefan Matile, and Aurélien Roux. A fluorescent membrane tension probe. Nature Chemistry, 10(11):1118–1125, 2018.

48. Henry De Belly, Shannon Yan, Hudson Borja da Rocha, Sacha Ichbiah, Jason P Town, Patrick J Zager, Dorothy C Estrada, Kirstin Meyer, Hervé Turlier, Carlos Bustamante, and Orion D Weiner. Cell protrusions and contractions generate long-range membrane tension propagation. Cell, 186(14):3049–3061.e15, jul 2023.

49. Tony Yeung, Penelope C. Georges, Lisa A. Flanagan, Beatrice Marg, Miguelina Ortiz, Makoto Funaki, Nastaran Zahir, Wenyu Ming, Valerie Weaver, and Paul A. Janmey. Effects of substrate stiffness on cell morphology, cytoskeletal structure, and adhesion. Cell Motility and the Cytoskeleton, 60(1):24–34, jan 2005.

50. Marion Ghibaudo, Alexandre Saez, Léa Trichet, Alain Xayaphoummine, Julien Browaeys, Pascal Silberzan, Axel Buguin, and Benoît Ladoux. Traction forces and rigidity sensing regulate cell functions. Soft Matter, 4(9):1836, 2008.

51. Sam Walcott and Sean X. Sun. A mechanical model of actin stress fiber formation and substrate elasticity sensing in adherent cells. Proceedings of the National Academy of Sciences of the United States of America, 107(17):7757–7762, 2010.

52. Aleksi Isomursu, Keun Young Park, Jay Hou, Bo Cheng, Mathilde Mathieu, Ghaidan A. Shamsan, Benjamin Fuller, Jesse Kasim, M. Mohsen Mahmoodi, Tian Jian Lu, Guy M. Genin, Feng Xu, Min Lin, Mark D. Distefano, Johanna Ivaska, and David J. Odde. Directed cell migration towards softer environments. Nature Materials, 21(September), 2022.

53. J M Scholey, K A Taylor, and J Kendrick-Jones. Regulation of non-muscle myosin assembly by calmodulin-dependent light chain kinase. Nature, 287(5779):233–5, sep 1980.

54. M Uehata, T Ishizaki, H Satoh, T Ono, T Kawahara, T Morishita, H Tamakawa, K Yamagami, J Inui, M Maekawa, and S Narumiya. Calcium sensitization of smooth muscle mediated by a Rho-associated protein kinase in hypertension. Nature, 389(6654):990–4, oct 1997.

55. Sotaro Sakurada, Noriko Takuwa, Naotoshi Sugimoto, Yu Wang, Minoru Seto, Yasuharu Sasaki, and Yoh Takuwa. Ca2+-dependent activation of Rho and Rho kinase in membrane depolarization-induced and receptor stimulation-induced vascular smooth muscle contraction. Circulation research, 93(6):548–56, sep 2003.

56. Tatsuya Miyamoto, Tsutomu Mochizuki, Hiroshi Nakagomi, Satoru Kira, Masaki Watanabe, Yasunori Takayama, Yoshiro Suzuki, Schuichi Koizumi, Masayuki Takeda, and Makoto Tominaga. Functional role for Piezo1 in stretch-evoked Ca2+ influx and ATP release in Urothelial cell cultures. Journal of Biological Chemistry, 289(23):16565–16575, 2014.

57. Stuart M Cahalan, Viktor Lukacs, Sanjeev S Ranade, Shu Chien, Michael Bandell, and Ardem Patapoutian. Piezo1 links mechanical forces to red blood cell volume. eLife, 4, may 2015.

58. Shilong Yang, Xinwen Miao, Steven Arnold, Boxuan Li, Alan T. Ly, Huan Wang, Matthew Wang, Xiangfu Guo, Medha M. Pathak, Wenting Zhao, Charles D. Cox, and Zheng Shi. Membrane curvature governs the distribution of Piezo1 in live cells. Nature Communications, 13(1):7467, dec 2022.

59. Mingxi Yao, Ajay Tijore, Delfine Cheng, Jinyuan Vero Li, Anushya Hariharan, Boris Martinac, Guy Tran Van Nhieu, Charles D. Cox, and Michael Sheetz. Force- and cell state–dependent recruitment of Piezo1 drives focal adhesion dynamics and calcium entry. Science Advances, 8(45):1–15, 2022.

60. Sheryl P. Denker, Derek C. Huang, John Orlowski, Heinz Furthmayr, and Diane L. Barber. Direct binding of the Na-H exchanger NHE1 to ERM proteins regulates the cortical cytoskeleton and cell shape independently of H+ translocation. Molecular Cell, 6(6):1425–1436, 2000.

61. Sheryl P Denker and Diane L Barber. Cell migration requires both ion translocation and cytoskeletal anchoring by the Na-H exchanger NHE1. The Journal of cell biology, 159(6):1087–96, dec 2002.

62. S Wakabayashi, T Ikeda, T Iwamoto, J Pouysségur, and M Shigekawa. Calmodulinbinding autoinhibitory domain controls “pH-sensing” in the Na+/H+ exchanger NHE1 through sequence-specific interaction. Biochemistry, 36(42):12854–61, oct 1997.

63. Sangkyun Cho, Jerome Irianto, and Dennis E Discher. Mechanosensing by the nucleus: From pathways to scaling relationships. The Journal of cell biology, 216(2):305–315, feb 2017.

64. Panagiotis Mistriotis, Emily O. Wisniewski, Kaustav Bera, Jeremy Keys, Yizeng Li, Soontorn Tuntithavornwat, Robert A. Law, Nicolas A. Perez-Gonzalez, Eda Erdogmus, Yuqi Zhang, Runchen Zhao, Sean X. Sun, Petr Kalab, Jan Lammerding, and Konstantinos Konstantopoulos. Confinement hinders motility by inducing RhoA-mediated nuclear influx, volume expansion, and blebbing. Journal of Cell Biology, 218(12):4093–4111, dec 2019.

65. A. J. Lomakin, C. J. Cattin, D. Cuvelier, Z. Alraies, M. Molina, G. P.F. Nader, N. Srivastava, P. J. Saez, J. M. Garcia-Arcos, I. Y. Zhitnyak, A. Bhargava, M. K. Driscoll, E. S. Welf, R. Fiolka, R. J. Petrie, N. S. de Silva, J. M. González-Granado, N. Manel, A. M. Lennon-Duménil, D. J. Müller, and M. Piel. The nucleus acts as a ruler tailoring cell responses to spatial constraints. Science, 370(6514), 2020.

66. Valeria Venturini, Fabio Pezzano, Frederic Català Castro, Hanna Maria Häkkinen, Senda Jiménez-Delgado, Mariona Colomer-Rosell, Monica Marro, Queralt Tolosa-Ramon, Sonia Paz-López, Miguel A. Valverde, Julian Weghuber, Pablo Loza-Alvarez, Michael Krieg, Stefan Wieser, and Verena Ruprecht. The nucleus measures shape changes for cellular proprioception to control dynamic cell behavior. Science, 370(6514), 2020.

67. Yohalie Kalukula, Andrew D. Stephens, Jan Lammerding, and Sylvain Gabriele. Mechanics and functional consequences of nuclear deformations. Nature Reviews Molecular Cell Biology, 23(9):583–602, sep 2022.

68. Yang Song, Jennifer Soto, Binru Chen, Tyler Hoffman, Weikang Zhao, Ninghao Zhu, Qin Peng, Longwei Liu, Chau Ly, Pak Kin Wong, Yingxiao Wang, Amy C Rowat, Siavash K Kurdistani, and Song Li. Transient nuclear deformation primes epigenetic state and promotes cell reprogramming. Nature Materials, 21(10):1191–1199, oct 2022.

69. Jerome Irianto, Joe Swift, Rui P. Martins, Graham D. McPhail, Martin M. Knight, Dennis E. Discher, and David A. Lee. Osmotic challenge drives rapid and reversible chromatin condensation in chondrocytes. Biophysical Journal, 104(4):759–769, 2013.

70. Andreas Krämer, Jeff Green, Jack Pollard, and Stuart Tugendreich. Causal analysis approaches in Ingenuity Pathway Analysis. Bioinformatics, 30(4):523–530, feb 2014.

71. Jordi Rius, Monica Guma, Christian Schachtrup, Katerina Akassoglou, Annelies S Zinkernagel, Victor Nizet, Randall S Johnson, Gabriel G Haddad, and Michael Karin. NF-kappaB links innate immunity to the hypoxic response through transcriptional regulation of HIF1alpha. Nature, 453(7196):807–11, jun 2008.

72. Kuppusamy Balamurugan, Ju-Ming Wang, Hsin-Hwa Tsai, Shikha Sharan, Miriam Anver, Robert Leighty, and Esta Sterneck. The tumour suppressor C/EBP*δ* inhibits FBXW7 expression and promotes mammary tumour metastasis. The EMBO journal, 29(24):4106–17, dec 2010.

73. Chuanzhao Zhang, Wanqing Iris Zhi, Haiquan Lu, Debangshu Samanta, Ivan Chen, Edward Gabrielson, and Gregg L Semenza. Hypoxia-inducible factors regulate pluripotency factor expression by ZNF217- and ALKBH5-mediated modulation of RNA methylation in breast cancer cells. Oncotarget, 7(40):64527–64542, oct 2016.

74. Der-Chih Yang, Muh-Hwa Yang, Chih-Chien Tsai, Tung-Fu Huang, Yau-Hung Chen, and Shih-Chieh Hung. Hypoxia inhibits osteogenesis in human mesenchymal stem cells through direct regulation of RUNX2 by TWIST. PloS one, 6(9):e23965, 2011.

75. Kay-Dietrich Wagner, Nicole Wagner, Sven Wellmann, Gunnar Schley, Anja Bondke, Heinz Theres, and Holger Scholz. Oxygen-regulated expression of the Wilms’ tumor suppressor Wt1 involves hypoxia-inducible factor-1 (HIF-1). FASEB journal : official publication of the Federation of American Societies for Experimental Biology, 17(10):1364–6, jul 2003.

76. Roland Lang and Faizal A M Raffi. Dual-Specificity Phosphatases in Immunity and Infection: An Update. International journal of molecular sciences, 20(11), jun 2019.

77. Yohannes Mebratu and Yohannes Tesfaigzi. How ERK1/2 activation controls cell proliferation and cell death: Is subcellular localization the answer? Cell cycle (Georgetown, Tex.), 8(8):1168–75, apr 2009.

78. Xing Chen, Shan Lin, Ying Lin, Songsong Wu, Minling Zhuo, Ailong Zhang, Junjie Zheng, and Zhenhui You. BRAF-activated WT1 contributes to cancer growth and regulates autophagy and apoptosis in papillary thyroid carcinoma. Journal of translational medicine, 20(1):79, feb 2022.

79. J Chung, E Uchida, T C Grammer, and J Blenis. STAT3 serine phosphorylation by ERK-dependent and -independent pathways negatively modulates its tyrosine phosphorylation. Molecular and cellular biology, 17(11):6508–16, nov 1997.

80. H Gille, M Kortenjann, O Thomae, C Moomaw, C Slaughter, M H Cobb, and P E Shaw. ERK phosphorylation potentiates Elk-1-mediated ternary complex formation and transactivation. The EMBO journal, 14(5):951–62, mar 1995.

81. Christopher L. Yankaskas, Keyata N. Thompson, Colin D. Paul, Michele I. Vitolo, Panagiotis Mistriotis, Ankit Mahendra, Vivek K. Bajpai, Daniel J. Shea, Kristen M. Manto, Andreas C. Chai, Navin Varadarajan, Aikaterini Kontrogianni-Konstantopoulos, Stuart S. Martin, and Konstantinos Konstantopoulos. A microfluidic assay for the quantification of the metastatic propensity of breast cancer specimens. Nature Biomedical Engineering, 3(6):452–465, 2019.

82. Cary R. Boyd-Shiwarski, Daniel J. Shiwarski, Shawn E. Griffiths, Rebecca T. Beacham, Logan Norrell, Daryl E. Morrison, Jun Wang, Jacob Mann, William Tennant, Eric N. Anderson, Jonathan Franks, Michael Calderon, Kelly A. Connolly, Muhammad Umar Cheema, Claire J. Weaver, Lubika J. Nkashama, Claire C. Weckerly, Katherine E. Querry, Udai Bhan Pandey, Christopher J. Donnelly, Dandan Sun, Aylin R. Rodan, and Arohan R. Subramanya. WNK kinases sense molecular crowding and rescue cell volume via phase separation. Cell, 185(24):4488–4506.e20, 2022.

83. Yan-Jun Guo, Wei-Wei Pan, Sheng-Bing Liu, Zhong-Fei Shen, Ying Xu, and Ling-Ling Hu. ERK/MAPK signalling pathway and tumorigenesis. Experimental and therapeutic medicine, 19(3):1997–2007, mar 2020.

84. David Shorthouse, Angela Riedel, Emma Kerr, Luisa Pedro, Dóra Bihary, Shamith Samarajiwa, Carla P. Martins, Jacqueline Shields, and Benjamin A. Hall. Exploring the role of stromal osmoregulation in cancer and disease using executable modelling. Nature Communications, 9(1), 2018.

85. Bo Yang, Haguy Wolfenson, Vin Yee Chung, Naotaka Nakazawa, Shuaimin Liu, Junqiang Hu, Ruby Yun Ju Huang, and Michael P. Sheetz. Stopping transformed cancer cell growth by rigidity sensing. Nature Materials, 19(2):239–250, 2020.

86. Ajay Tijore, Mingxi Yao, Yu Hsiu Wang, Anushya Hariharan, Yasaman Nematbakhsh, Bryant Lee Doss, Chwee Teck Lim, and Michael Sheetz. Selective killing of transformed cells by mechanical stretch. Biomaterials, 275(May), 2021.

87. Rachelle N. Palchesko, Ling Zhang, Yan Sun, and Adam W. Feinberg. Development of Poly-dimethylsiloxane Substrates with Tunable Elastic Modulus to Study Cell Mechanobiology in Muscle and Nerve. PLoS ONE, 7(12):e51499, dec 2012.

