## Supplementary Text for "Cytoskeletal activation of NHE1 regulates mechanosensitive cell volume adaptation and proliferation"

Qin Ni,[1,2,†] Zhuoxu Ge,[1,2,†] Yizeng Li,[3], Gabriel Shatkin[4], Jinyu Fu,[1,5], Kaustav Bera,[1,6], Yuhang Yang[7], Yichen Wang [2], Anindya Sen[1,6], Yufei Wu[1,2], Ana Carina Nogueira Vasconcelos [1,2], Andrew P. Feinberg[4,7,8], Konstantinos Konstantopoulos,[1,4,6,7] Sean X. Sun [1,2,*]

[1]Institute for NanoBioTechnology, Johns Hopkins University, Baltimore, MD, USA

[2]Department of Mechanical Engineering, Johns Hopkins University, Baltimore, MD, USA

[3]Department of Biomedical Engineering, Binghamton University, Binghamton, NY, USA

[4]Department of Biomedical Engineering, Johns Hopkins University, Baltimore, MD, USA

[5]Department of Physics, Johns Hopkins University, Baltimore, MD, USA

[6]Department of Chemical and Biomolecular Engineering, Johns Hopkins University, Baltimore, MD, USA

[7]Department of Oncology, The Sidney Kimmel Comprehensive Cancer Center, Johns Hopkins University School of Medicine, Baltimore, USA

[8]Center for Epigenetics, Johns Hopkins University School of Medicine, Baltimore, MD, USA

[†]These authors contribute equally

#### 1 A comprehensive search of ion transporter expression in 3T3.

To obtain the RNA expression of all ion transporters in 3T3, we created an ion transporter list based on Transporter Classification Database (TCDB) (*1*) and the HUGO Gene Nomenclature Committee (HGNC) database (*2*). The list that contains 665 ion transporters can be found in supplementary files. We performed RNA seq on 3T3 and HT1080 in isotonic condition as discussed in the main text and plotted all ion transporters that express at least 10 transcripts per million (TPM) reads (Fig. ST-1). We categorize ion transporters into four large categories: (1) Passive ion channels including sodium channels, potassium channels, anion channels, calcium channels, transient receptor potential (TRP) channels, and ligand-gated ion channels. (2) ATPases. (3) Solute carrier (SLC) family. (4) Other ion related transporters including connexins, pannexins, and aquaporins.

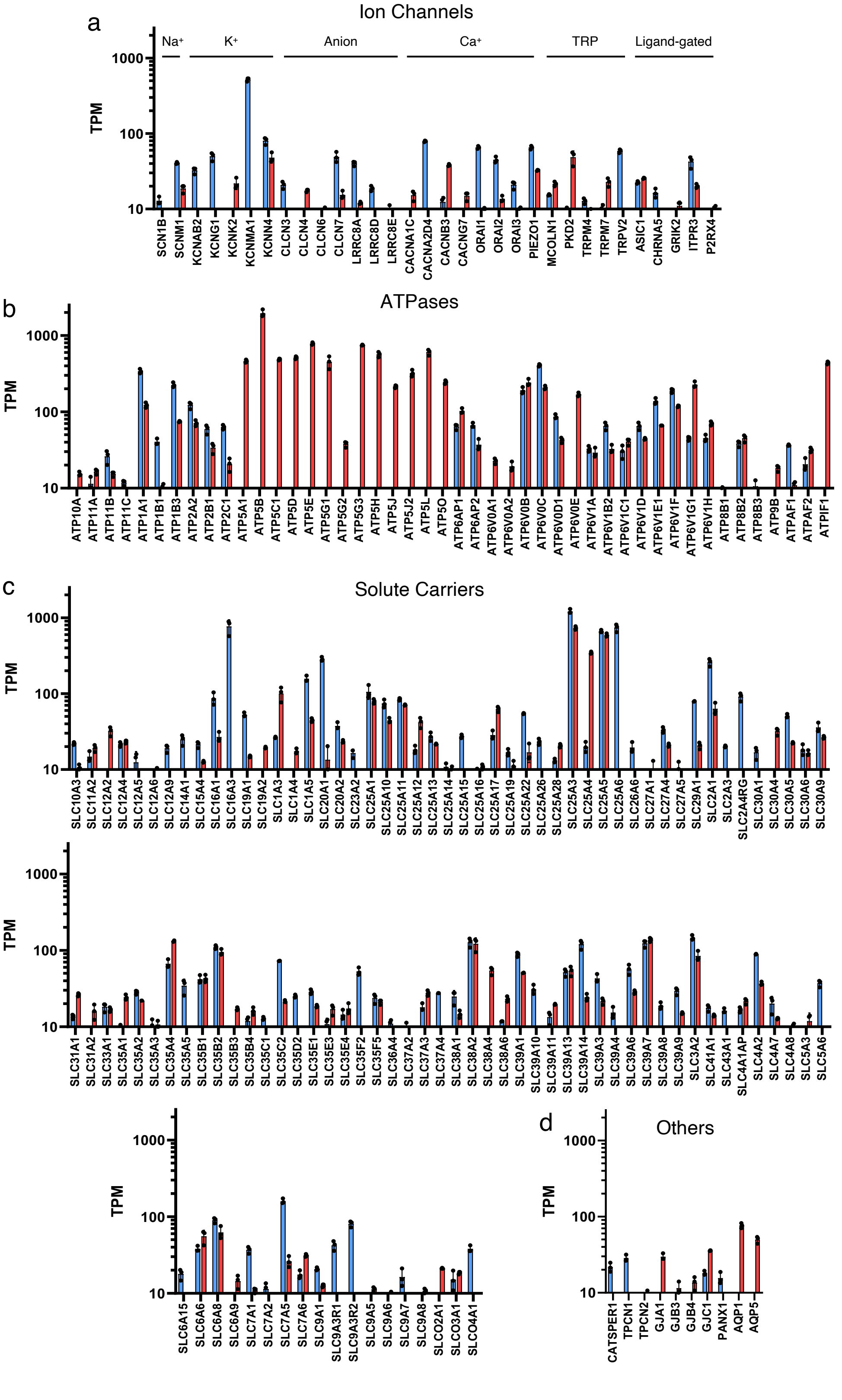


Figure ST-1: **RNA seq reveals ion transporters expression in 3T3 and HT1080.**

Figure S1: **RNA seq reveals ion transporters expression in 3T3 and HT1080.** (a-d) Ion transporters expressed at least 10 transcripts per million (TPM) in either 3T3 (red) or HT1080 (blue). Ion transporters are characterized into (a) passive ion channels (b) ATPases, (c) solute carriers, and (d) others. N = 3 biological repeats. Error bar represents standard deviation.

### 2 Model Description

#### 2.1 Model overview

The model of cell volume considers the coupled interactions between different ion transporters and the cell mechanics. The model details can be found in Ref. 3. Briefly, we consider the following intracellular species: $\mathrm{Na}^+, \mathrm{K}^+, \mathrm{Cl}^-, \mathrm{H}^+, \mathrm{HCO}_3^-, \mathrm{A}^-, \mathrm{Buf}^-$, and HBuf. $\mathrm{A}^-$ is the charged organic molecules and proteins that are not permeable across the cell membrane. $\mathrm{Buf}^-$ and HBuf are unprotonated buffer and protonated buffer species, respectively, where the total buffer $\mathrm{Buf}^- + \mathrm{HBuf}$ concentration is fixed and both species are non-permeable across the cell membrane. The cell is assumed to be spherical, and the change of cell radius is controlled by water flux across the cell surface ($J_{\mathrm{water}}$) as

$$\frac{dr}{dt} = J_{\mathrm{water}} = -\alpha_W(\Delta p - \Delta\Pi). \tag{ST1}$$

In this equation, the water flux is determined by the difference between the hydrostatic pressure ($\Delta p$) and the osmotic pressure gradient ($\Delta\Pi$) across the cell surface (*4, 5*). $\alpha_W$ is the combined permeability coefficient of water of cell lipid membrane and aquaporins. The force generated by hydrostatic pressure on cell surface is under force balance with cell membrane and the actomyosin cortical tension, which can be written by Laplace Law as $\Delta p = 2h\sigma/r$. The osmotic pressure gradient is written as

$$\Delta\Pi = RT(\sum_n c_n - \sum_n c_n^0), \tag{ST2}$$

where $c_n = \{c_{\mathrm{Na}}, c_{\mathrm{K}}, c_{\mathrm{Cl}}, c_{\mathrm{H}}, c_{\mathrm{HCO_3}}, c_{\mathrm{A}}, c_{\mathrm{Buf}}, c_{\mathrm{HBuf}}\}^T$ is the intracellular osmolarity, and $c_n^0 = \{c^0_{\mathrm{Na}}, c^0_{\mathrm{K}}, c^0_{\mathrm{Cl}}, c^0_{\mathrm{H}}, c^0_{\mathrm{HCO_3}}, c^0_{\mathrm{A}}, c^0_{\mathrm{Buf}}, c^0_{\mathrm{HBuf}}\}^T$ is the extracellular osmolarity.

We treated $\mathrm{Na}^+, \mathrm{K}^+$, and $\mathrm{Cl}^-$ as non-reactive ion species, and the their conservation is written as

$$\frac{d}{dt}(Vc_n) = 4\pi r^2 J_n, n \in \{\mathrm{Na}^+, \mathrm{K}^+, \mathrm{Cl}^-\}, \tag{ST3}$$

where $J_n$ is the net ion flux across the cell membrane for each species through active and passive ion transporters, and $V = 4/3\pi r^3$ is the cell volume. The conservation equation for the intracellular reactive species is

$$\frac{d}{dt}(Vc_{\mathrm{H}}) - 4\pi r^2 J_{\mathrm{H}} = \frac{d}{dt}(Vc_{\mathrm{HCO_3}}) - 4\pi r^2 J_{\mathrm{HCO_3}} + \frac{d}{dt}(Vc_{\mathrm{Buf}}). \tag{ST4}$$

We considered the following ion transporters: passive $\mathrm{Na}^+, \mathrm{K}^+$, and $\mathrm{Cl}^-$ channels, $\mathrm{Na}^+$-$\mathrm{K}^+$-$\mathrm{Cl}^-$ co-transporter (NKCC), $\mathrm{Na}^+$-$\mathrm{K}^+$ ATPase (NKA), $\mathrm{Na}^+$-$\mathrm{H}^+$ exchanger (NHE), $\mathrm{Na}^+$-$\mathrm{HCO}_3^-$ co-transporter (NBC), and anion exchange protein 2 (AE2). The net flux for each ion species is modeled as the sum of fluxes through the relevant transporters:

$$J_{\mathrm{Na}} = J_{\mathrm{Na},p} + J_{\mathrm{NKCC,Na}} + J_{\mathrm{NKA,Na}} + J_{\mathrm{NHE,Na}} + J_{\mathrm{NBC,Na}}, \tag{ST5}$$

$$J_{\mathrm{K}} = J_{\mathrm{K},p} + J_{\mathrm{NKCC,K}} + J_{\mathrm{NKA,K}}, \tag{ST6}$$

$$J_{\mathrm{Cl}} = J_{\mathrm{Cl},p} + J_{\mathrm{NKCC,Cl}} + J_{\mathrm{AE2,Cl}}, \tag{ST7}$$

$$J_{\mathrm{H}} = J_{\mathrm{NHE,H}}, \tag{ST8}$$

$$J_{\mathrm{HCO_3}} = J_{\mathrm{AE2,HCO_3}} + J_{\mathrm{NBC,HCO_3}}. \tag{ST9}$$

Passive ion transporters are typically gated by the membrane electrochemical potential or the membrane tension, and thus we modeled the passive ion fluxes as

$$J_{n,p} = \alpha_{n,p} G_m (RT \ln\Gamma_n - z_n F V_m), n \in \{\mathrm{Na}^+, \mathrm{K}^+, \mathrm{Cl}^-\} \tag{ST10}$$

where $V_m$ is the membrane potential, $z_n$ is the valence of each ion species, $\Gamma_n = c_n^0/c_n$ is the ratio of extra- to intra- cellular ion concentrations, $\alpha_{n,p}$ is the permeability coefficient of each species, and $G_m = [1 + e^{-\beta_1(\tau_m - \beta_2)}]^{-1} \in \{0, 1\}$ is a mechanosensitive gating function. NKCC flux is modeled as

$$J_{\mathrm{NKCC}} = J_{\mathrm{NKCC,Na}} = J_{\mathrm{NKCC,K}} = \frac{1}{2} J_{\mathrm{NKCC,Cl}} = \alpha_{\mathrm{NKCC}} RT (\ln\Gamma_{\mathrm{Na}} + \ln\Gamma_{\mathrm{K}} + 2\ln\Gamma_{\mathrm{Cl}}), \tag{ST11}$$

where $\alpha_{\mathrm{NKCC}}$ is the permeability coefficient independent of the cortical tension. NHE is modeled as a pH gated function:

$$J_{\mathrm{NHE}} = J_{\mathrm{NHE,Na}} = -J_{\mathrm{NHE,H}} = \alpha_{\mathrm{NHE}} G_{\mathrm{NHE}} RT (\ln\Gamma_{\mathrm{Na}} - \ln\Gamma_{\mathrm{H}}), \tag{ST12}$$

where $\alpha_{\mathrm{NHE}}$ is the permeability coefficient of NHE, which is assumed as a constant. $G_{\mathrm{NHE}} = [1 + e^{\beta_{\mathrm{NHE},1}(\mathrm{pH} - \beta_{\mathrm{NHE},2})}]^{-1}$ is a pH-gated function of NHE. Ion fluxes through AE2 takes a similar form:

$$J_{\mathrm{AE2}} = J_{\mathrm{AE2,Cl}} = -J_{\mathrm{AE2,HCO_3}} = \alpha_{\mathrm{AE2}} G_{\mathrm{AE2}} RT (\ln\Gamma_{\mathrm{Cl}} - \ln\Gamma_{\mathrm{HCO_3}}), \tag{ST13}$$

where $\alpha_{\mathrm{AE2}}$ is the permeability coefficient of AE2 that is assumed to be a constant, and $G_{\mathrm{AE2}} = [1 + e^{-\beta_{\mathrm{AE2},1}(\mathrm{pH} - \beta_{\mathrm{AE2},2})}]^{-1}$ is the pH-gated function of AE2. Ion fluxes through NBC depends on the electrochemical potential of $\mathrm{Na}^+$ and $\mathrm{HCO}_3$, which is modeled as

$$J_{\mathrm{NBC}} = J_{\mathrm{NBC,Na}} = -\frac{3}{2} J_{\mathrm{NBC,HCO_3}} = \alpha_{\mathrm{NBC}} [RT \ln(\Gamma_{\mathrm{Na}} \Gamma_{\mathrm{HCO_3}}^2) - (z_{\mathrm{Na}} + 2 z_{\mathrm{HCO_3}}) F V_m], \tag{ST14}$$

where $\alpha_{\mathrm{NBC}}$ is the permeability coefficient of NBC. Ion fluxes through NKA is modeled as

$$J_{\mathrm{NKA}} = J_{\mathrm{NKA,Na}} = -\frac{3}{2}J_{\mathrm{NKA,K}} = -\alpha_{\mathrm{NKA}}G_{V,\mathrm{NKA}}(1+\beta_{\mathrm{NKA,Na}}\Gamma_{\mathrm{Na}})^{-3}(1+\beta_{\mathrm{NKA,K}}/\Gamma_{\mathrm{K}})^{-2}, \tag{ST15}$$

where $\alpha_{\mathrm{NKA}}$ is the permeability coefficient of NKA, $\beta_{\mathrm{NKA,Na}}$ and $\beta_{\mathrm{NKA,K}}$ are scaling constant for $\Gamma_{\mathrm{Na}}$ and $\Gamma_{\mathrm{K}}$, respectively. $G_{V,\mathrm{NKA}} = 2[1 + e^{-\beta_{\mathrm{NKA},1}(V_m - \beta_{\mathrm{NKA},2})}]^{-1}$ is the voltage-gated activity function of NKA.

Inside the cell, we modeled the bicarbonate-carbonic acid equilibrium as

$$\mathrm{CO_2(aq)} + \mathrm{H_2O(l)} \rightleftharpoons \mathrm{H^+(aq)} + \mathrm{HCO_3^-(aq)},$$

where $[\mathrm{CO_2}]_{\mathrm{aq}}$ depends on the partial pressure of $CO_2$, $P_{\mathrm{CO_2}}$, by Henry's constant, $k_H$, i.e. $[\mathrm{CO_2}]_{\mathrm{aq}} = P_{\mathrm{CO_2}}/k_H$. For intracellular pH, we have

$$\mathrm{pH} - \mathrm{p}K_c = \log_{10}\frac{[\mathrm{HCO_3^-}]_{\mathrm{aq}}}{[\mathrm{CO_2}]_{\mathrm{aq}}},$$

where $\mathrm{pH} = -\log_{10}[\mathrm{H^+}]_{\mathrm{aq}}$ is the intracellular pH, $K_c = [\mathrm{HCO_3^-}]_{\mathrm{aq}}[\mathrm{H^+}]_{\mathrm{aq}}/[\mathrm{CO_2}]_{\mathrm{aq}}$ is the equilibrium constant for the bicarbonate-carbonic acid pair, and $\mathrm{p}K_c = \log_{10}K_c$. Cells also contain various buffer solutions, and their reactions can be written as

$$\mathrm{H_\ell Buf(aq)} \rightleftharpoons \ell\mathrm{H^+(aq)} + \mathrm{Buf^{\ell-}(aq)},$$

where $\ell$ = 1, 2, 3, ... for different buffer species. $K_{B,\ell} = [\mathrm{Buf^{\ell-}}]_{\mathrm{aq}}[\mathrm{H^+}]_{\mathrm{aq}}^{\ell}/[\mathrm{H_\ell Buf}]_{\mathrm{aq}}$ is the equilibrium constant, and $\mathrm{p}K_{B,\ell} = \log_{10}K_{B,\ell}$. We set $\ell$ = 1 by default.We also assume both the intracellular and the extracellular are electro-neutral, which is realized by enforcing $\sum z_n c_n = 0$.

Overall, the full set of equations is as follows:

$$\frac{dr}{dt} = J_{\mathrm{water}} = -\alpha_W\left[\frac{2h\sigma}{r} - RT\left(\sum c_n - \sum c_n^0\right)\right], \tag{ST16}$$

$$\sum z_n c_n = 0, \tag{ST17}$$

$$\frac{d}{dt}(Vc_{\mathrm{Na}}) = 4\pi r^2(J_{\mathrm{Na},p} + J_{\mathrm{NKCC}} + J_{\mathrm{NKA}} + J_{\mathrm{NHE}} + J_{\mathrm{NBC}}), \tag{ST18}$$

$$\frac{d}{dt}(Vc_{\mathrm{K}}) = 4\pi r^2(J_{\mathrm{K},p} + J_{\mathrm{NKCC}} - \frac{2}{3}J_{\mathrm{NKA}}), \tag{ST19}$$

$$\frac{d}{dt}(Vc_{\mathrm{Cl}}) = 4\pi r^2(J_{\mathrm{Cl},p} + 2J_{\mathrm{NKCC}} + J_{\mathrm{AE2}}), \tag{ST20}$$

$$\frac{d}{dt}(Vc_{\mathrm{H}}) + 4\pi r^2 J_{\mathrm{NHE}} = \frac{d}{dt}(Vc_{\mathrm{HCO_3}}) + 4\pi r^2(J_{\mathrm{AE2}} - n_{\mathrm{NBC}}J_{\mathrm{NBC}}) + \frac{d}{dt}(Vc_{\mathrm{Buf}}), \tag{ST21}$$

Table ST-1: Generic model parameters.

| Parameter | Description | Value | Source |
|---|---|---|---|
| $h$ (nm) | Cortical thickness | 500 | (*6*) |
| $R$ (J/mol/K) | Ideal gas constant | 8.31 | Physical constant |
| $T$ (K) | Absolute temperature | 310 | Physiological condition |
| $\sigma_a$ (Pa) | Active contraction | $10^3$ | (*6*) |
| $N_A$ (mol) | Total intracellular $A^-$ | 0.142 | Estimated |
| $N_{\mathrm{Bul}} + N_{\mathrm{HBul}}$ (pmol) | Total intracellular Buffer Solution | 0.16 | Estimated |
| $\alpha_{\mathrm{w}}$ (m/Pa/s) | Permeability coefficient of water | $10^{-10}$ | (*4*) |
| $\beta_{\mathrm{NKA,Na}}$ | Constant in $J_{\mathrm{NKA}}$ | 0.1 | Estimated |
| $\beta_{\mathrm{NKA,K}}$ | Constant in $J_{\mathrm{NKA}}$ | 10 | Estimated |
| $\beta_1$ (m/N) | In $G_m = [1 + e^{-\beta_1(\tau_m - \beta_2)}]^{-1}$ | $2 \times 10^3$ | Estimated |
| $\beta_2$ (N/m) | In $G_m = [1 + e^{-\beta_1(\tau_m - \beta_2)}]^{-1}$ | $5 \times 10^{-4}$ | Estimated |
| $\beta_{\mathrm{NKA,1}}$ (1/mV) | In $G_{V,\mathrm{NKA}} = 2/[1 + e^{-\beta_{\mathrm{NKA,1}}(V_m - \beta_{\mathrm{NKA,2}})}] - 1$ | 0.03 | (*7*) |
| $\beta_{\mathrm{NKA,2}}$ (mV) | In $G_{V,\mathrm{NKA}} = 2/[1 + e^{-\beta_{\mathrm{NKA,1}}(V_m - \beta_{\mathrm{NKA,2}})}] - 1$ | $-150$ | (*7*) |
| $\beta_{\mathrm{NHE,1}}$ | In $G_{\mathrm{NHE}} = [1 + e^{\beta_{\mathrm{NHE,1}}(\mathrm{pH} - \beta_{\mathrm{NHE,2}})}]^{-1}$ | 15 | (*8*) |
| $\beta_{\mathrm{NHE,2}}$ | In $G_{\mathrm{NHE}} = [1 + e^{\beta_{\mathrm{NHE,1}}(\mathrm{pH} - \beta_{\mathrm{NHE,2}})}]^{-1}$ | 7.2 | (*8*) |
| $\beta_{AE,1}$ | In $G_{\mathrm{AE2}} = [1 + e^{-\beta_{AE,1}(\mathrm{pH} - \beta_{AE,2})}]^{-1}$ | 10 | (*8*) |
| $\beta_{AE,2}$ | In $G_{\mathrm{AE2}} = [1 + e^{-\beta_{AE,1}(\mathrm{pH} - \beta_{AE,2})}]^{-1}$ | 7.1 | (*8*) |
| $k_H$ (atm/M) | Henry's constant | 29 | (*9*) |
| $P_{\mathrm{CO_2}}$ (atm) | Partial pressure of $CO_2$ | 5% | Physiological condition |
| $K_c$ | p$K$ for bicarbonate-carbonic acid pair | 6.1 | (*9*) |
| $K_B$ | p$K$ for intracellular buffer | 6.7 | Estimated |
| $c^0_{\mathrm{Na}}$ (mM) | $Na^+$ concentration in the medium | 145 | Physiological condition |
| $c^0_{\mathrm{K}}$ (mM) | $K^+$ concentration in the medium | 9 | Physiological condition |
| $c^0_{\mathrm{Cl}}$ (mM) | $Cl^-$ concentration in the medium | 105 | Physiological condition |
| $c^0_{\mathrm{HCO_3}}$ (mM) | $HCO_3^-$ concentration in the medium | 35 | Physiological condition |
| $c^0_{\mathrm{G}}$ (mM) | Molecular concentration in the medium | 25 | Physiological condition |

In this model, we assume $c^0_{\mathrm{Na}}, c^0_{\mathrm{K}}, c^0_{\mathrm{Cl}}, c^0_{\mathrm{H}}, c^0_{\mathrm{HCO_3}}, N_{\mathrm{A}}, P_{\mathrm{CO_2}}$, and $N_{\mathrm{HBuf}} + N_{\mathrm{Buf}}$ are known based the composition of cell culture media and $CO_2$. We solved 6 unknowns, $r, V_m, c_{\mathrm{Na}}, c_{\mathrm{K}}, c_{\mathrm{Cl}}$ and pH from Eq. 16-21. Other parameters are derived either from give quantities or from the unknowns.

All systems were initialized at the same condition, then we allowed each system to evolve to a steady state. The generic parameters that applicable to all cell types used in this work can found in Tab. ST-1. The hypotonic shock was applied by reducing 50% of the extracellular species concentration. The inhibition of NHE and NKA was conducted by reducing their permeability coefficients by 90%.

#### 2.2 Ion Transporter Activities and Automatic Volume Recovery

The expression and activity of membrane ion transporters vary among different cell types, and the response of cells to osmotic shocks depends on their overall ion transporter activities (*3*) .For example, from our experimental data, HT1080 and 3T3 cells display different levels of volume recovery after a hypotonic shock. However, in some cases, no volume recovery is observed unless parameters are adjusted following the application of a hypotonic shock (*3*).

To systematically investigate the impact of ion transporter activity on cell volume regulation during hypotonic shock, we employed a Monte Carlo search of widely-reported volume-regulated transporters, including NHE, NKCC, NKA, and passive Na, Cl, and K channels. Each parameter was randomly varied between $10^{-2}$ and $10^{2}$ times its baseline value, where the baseline value represents the typical level observed in a mammalian tissue cell exhibiting stable homeostasis. The baseline values for each transporter are listed in Tab. ST-2, where $P_0 = 3 \times 10^{-9}$ mol²⁄(J·m²·s) is the overall scaling factor. The rest of the parameters are provided in Tab. ST-1.

Using our experimental findings, we first identified ion transporter activities that result in (50±5)% cell volume recovery after hypotonic shock, as shown in Fig. ST-2a. The activity level of each transporter is normalized to its corresponding baseline value. We then conducted a principle component analysis (PCA) to examine the relative importance of ion transporters in controlling regulatory volume recovery. The first 6 principle components (PCs) explains the majority of the variance (Fig. ST-2b), indicating that ion transporter activities that mainly project onto these PCs are potentially less important since they do not need to be kept within a specific range. Therefore, we focused on the last 3 PCs and plotted the eigenvectors of the covariance matrix of all six ion transporters in Fig. ST-2c. We found that among active ion transporters, PCs 7-9 are mainly contributed by NHE and NKA. This observation suggests that the activities of NHE and NKA are two key parameters for regulatory volume recovery, while NKCC, NBC, and AE2 plays a less significant role. This finding is in agreement with our experimental observations. Among passive ion channels, the Na channels is the major contributor to PCs 7-9. We also plotted the projections of ion transporter parameters onto PCs 7-9 (Fig. ST-2d and e).

#### 2.3 HT1080 and 3T3 Cells

Using our experimental data, we have identified sets of membrane ion transporter activities that are representative of HT1080 and 3T3 cells. These parameters, which are derived from random simulations, are listed in Table ST-3, along with the remaining parameters provided in Table ST-

Table ST-2: Base-line activity values for each channel and pump.

| Parameter | Description | Value |
|---|---|---|
| $\alpha_{\text{Na},p}$ (mol$^2$/J/m$^2$/s) | Permeability coefficient of Na channel | $0.1P_0$ |
| $\alpha_{\text{K},p}$ (mol$^2$/J/m$^2$/s) | Permeability coefficient of K channel | $P_0$ |
| $\alpha_{\text{Cl},p}$ (mol$^2$/J/m$^2$/s) | Permeability coefficient of Cl channel | $0.2P_0$ |
| $\alpha_{\text{NKA}}$ (mol/m$^2$/s) | Permeability coefficient of NKA | $10^5P_0/RT$ |
| $\alpha_{\text{NKCC}}$ (mol$^2$/J/m$^2$/s) | Permeability coefficient of NKCC | $10^{-3}P_0$ |
| $\alpha_{\text{NHE}}$ (mol$^2$/J/m$^2$/s) | Permeability coefficient of NHE | $P_0$ |
| $\alpha_{\text{AE2}}$ (mol$^2$/J/m$^2$/s) | Permeability coefficient of AE2 | $0.1P_0$ |
| $\alpha_{\text{NBC}}$ (mol$^2$/J/m$^2$/s) | Permeability coefficient of NBC | $1.5P_0$ |

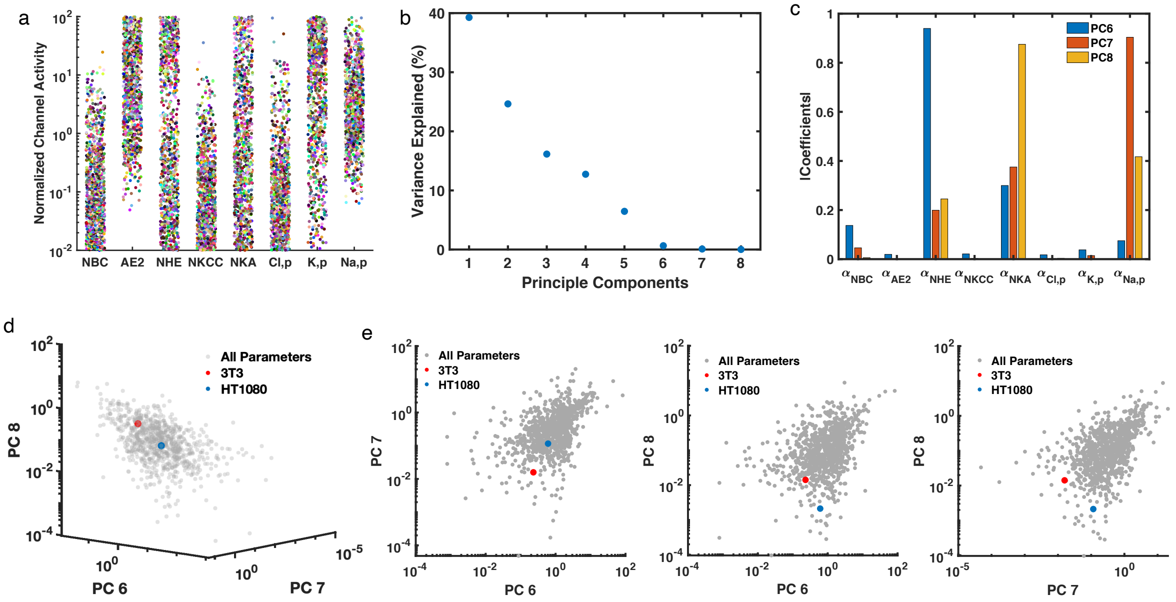


Figure ST-2: (a) Distribution of ion transporter activities that generate $(50 \pm 5)\%$ regulatory cell volume recovery upon a hypotonic shock. Each activity level is normalized with respect to the corresponding base-line value. The base-line values become 1 after normalization. Each color represents a set of parameters obtained from the parameter search. (b) Variance explained at each principle components (PCs) of all parameters presenting in (a). (c) Eigenvectors of the covariance matrix of parameters presenting in (a) on PC 7, 8, and 9. (d) All parameters (gray), and parameters of 3T3 (red) and HT1080 (blue) projected on PC 7, 8, and 9. (e) 2D projections of (d).

Table ST-3: Parameters for HT1080 and 3T3 cells.

| Parameter | Description | HT1080 | 3T3 |
|---|---|---|---|
| $\alpha_{\text{Na},p}$ (mol$^2$/J/m$^2$/s) | Permeability coefficient of Na channel | $1.12 \times 10^{-11}$ | $1.41 \times 10^{-11}$ |
| $\alpha_{\text{K},p}$ (mol$^2$/J/m$^2$/s) | Permeability coefficient of K channel | $1.27 \times 10^{-8}$ | $9.12 \times 10^{-10}$ |
| $\alpha_{\text{Cl},p}$ (mol$^2$/J/m$^2$/s) | Permeability coefficient of Cl channel | $8.78 \times 10^{-12}$ | $2.26 \times 10^{-8}$ |
| $\alpha_{\text{NKA}}$ (mol/m$^2$/s) | Permeability coefficient of NKA | $7.99 \times 10^{-9}$ | $3.06 \times 10^{-9}$ |
| $\alpha_{\text{NKCC}}$ (mol$^2$/J/m$^2$/s) | Permeability coefficient of NKCC | $2.04 \times 10^{-10}$ | $1.68 \times 10^{-12}$ |
| $\alpha_{\text{NHE}}$ (mol$^2$/J/m$^2$/s) | Permeability coefficient of NHE | $1.04 \times 10^{-9}$ | $1.19 \times 10^{-10}$ |
| $\alpha_{\text{AE2}}$ (mol$^2$/J/m$^2$/s) | Permeability coefficient of AE2 | $5.75 \times 10^{-12}$ | $1.73 \times 10^{-9}$ |
| $\alpha_{\text{NBC}}$ (mol$^2$/J/m$^2$/s) | Permeability coefficient of NBC | $5.03 \times 10^{-10}$ | $3.46 \times 10^{-8}$ |
| $K_B$ | p$K$ for intracellular buffer | 8.55 | 4.73 |

1. Their projections on PCs 4-6 are highlighted in Fig. ST-2d-e. It is important to note that the parameters in Table ST-3 represent the activity levels of the ion transporters, rather than their RNA expression levels. The activity levels are function of ion transporter expression as well as their activation by voltages, tension, and $[Ca^{2+}]$. These parameters are used to model the changes in cell volume following hypotonic shocks as depicted in the main text.

#### 2.4 Secondary volume increase in 3T3 Cells

The volume response of 3T3 cells exhibits a secondary volume increase (SVI) after a hypotonic shock. Based on our modeling, it is not possible to directly produce the SVI by solely adjusting parameters. This suggests the existence of additional activation mechanisms contributing to the SVI. As discussed in the main text, we attribute the additional volume increase to the activation of NHE following the initial volume recovery in response to the hypotonic shock. To incorporate this phenomenon into the model, we artificially introduced an NHE activation function in the 3T3 cells by prescribing an amplitude adjustment factor, $A(t)$, to the NHE flux expression in Eq. ST12.

This normalized factor is chosen by

$$A\left(t\right) = 1 + \gamma_1 \left(t - t_a\right) e^{-\gamma_2 t}, \tag{ST22}$$

where $\gamma_1$ and $\gamma_2$ are constant that modulate the amplitude of the SVI. We set $\gamma_1 = 1$ s$^{-1}$ and $\gamma_2 = 1 \times 10^{-3}$ s$^{-1}$ as a reference value based on the experimental data. The parameter $t_a$ represents the activation time of the SVI. The additional NHE activation is triggered at the same time as the hypotonic shock, as we assume that NHE is activated as the cell swells in response to the shock.

This activation factor is only applied to the control and NHE inhibition results; it is not included in the NKA inhibition results.

### 3 Statistics of time series plots

Here we present the statistical analysis of the time series data shown in the main and supplemental figures. The aim of this analysis is to establish the statistical significance of variations observed in the time series data across different experimental scenarios: between two model cell lines, between vehicle control and drug treatment, and between a genetically edited cell line and its control. All individual trajectories were initially normalized to their respective values prior to osmotic shock or mechanical stretch. Subsequently, Mann-Whitney U tests were performed to compare the two conditions at each time point following osmotic shock or mechanical stretch. The resulting p-values, plotted as a function of time, are shown in Figures ST-3 to ST-5.

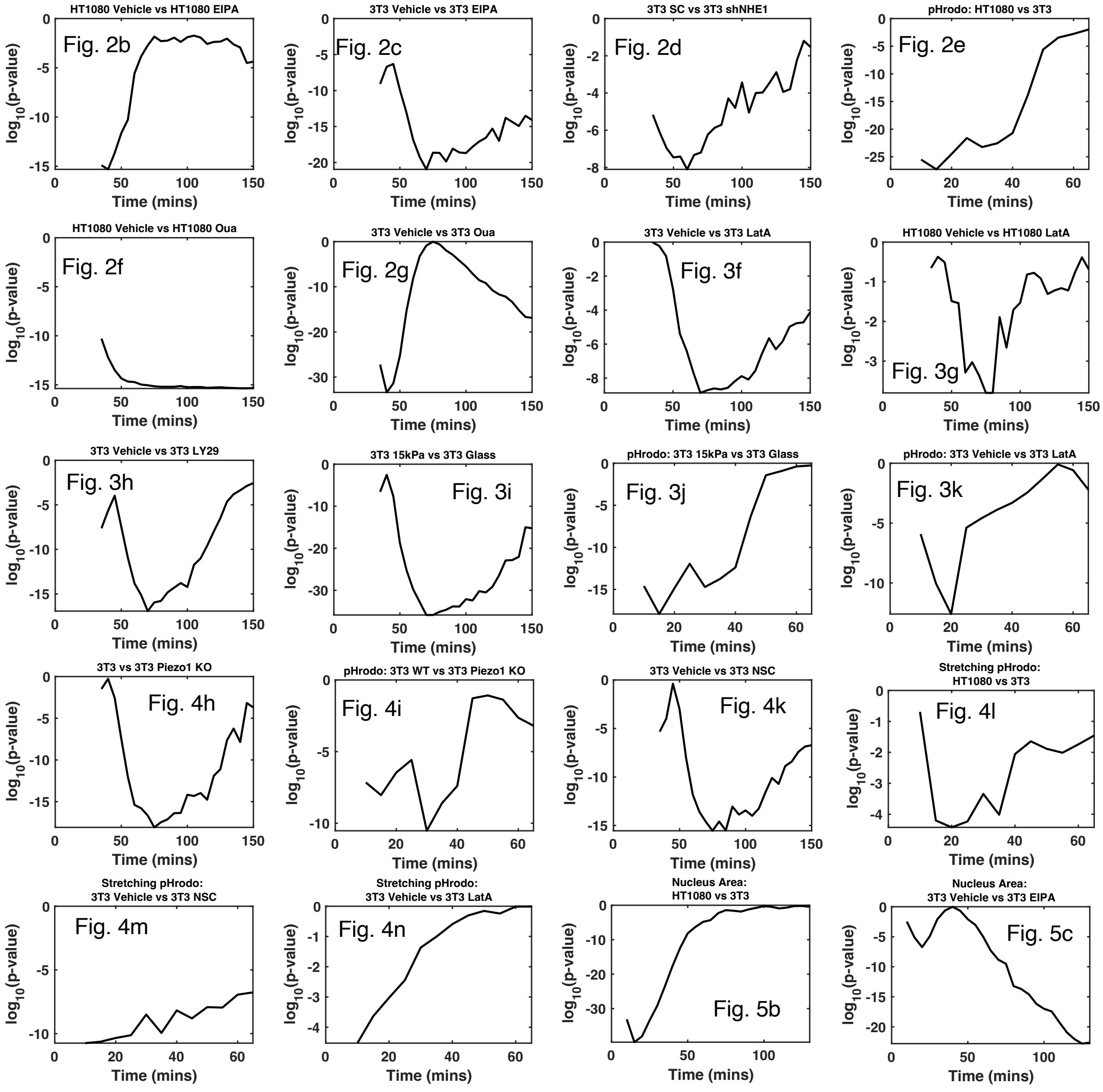


Figure ST-3: Statistical analysis of all time series plots in main figures.

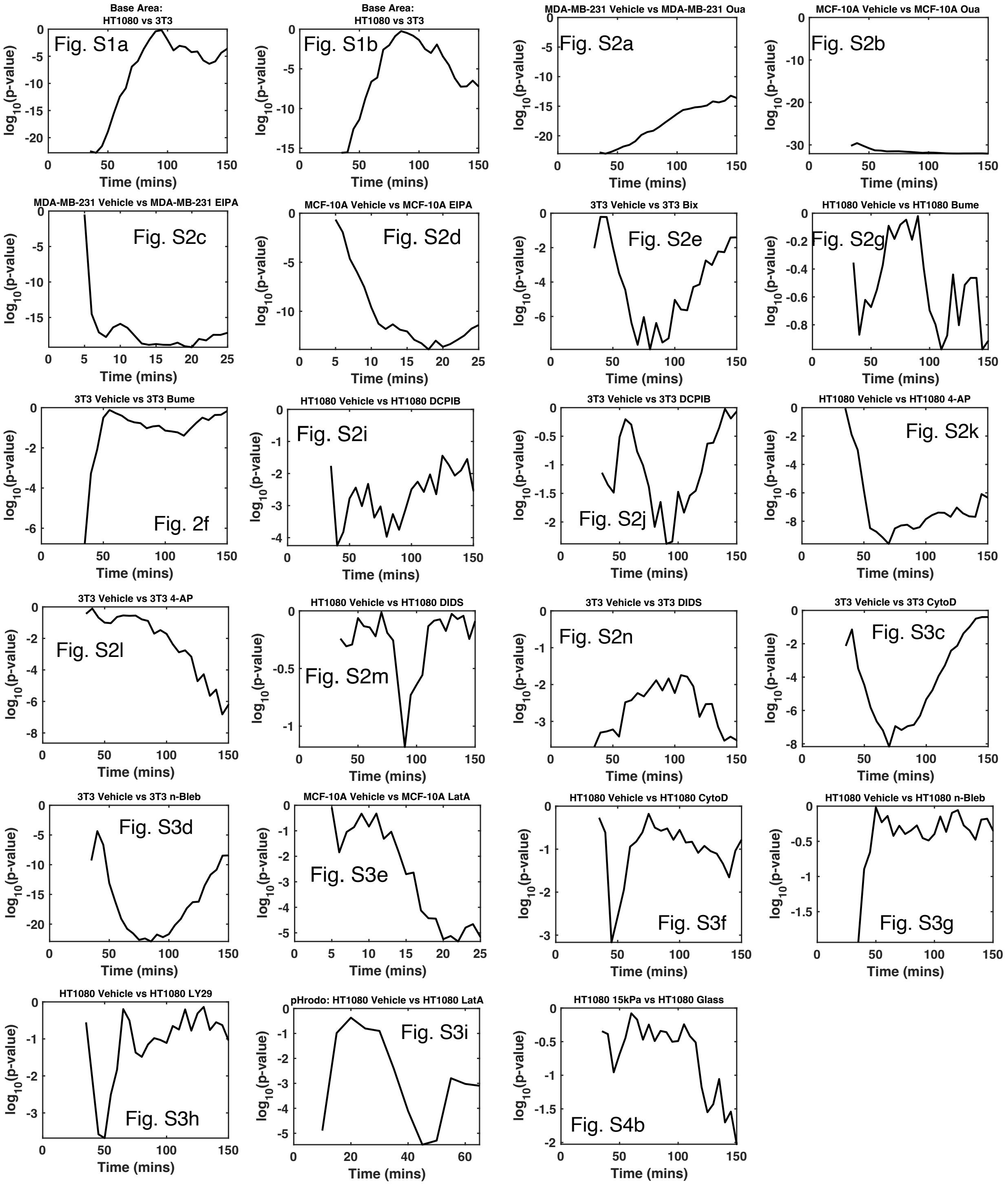


Figure ST-4: Statistical analysis of all time series plots in Supplementary figures S1-S4.

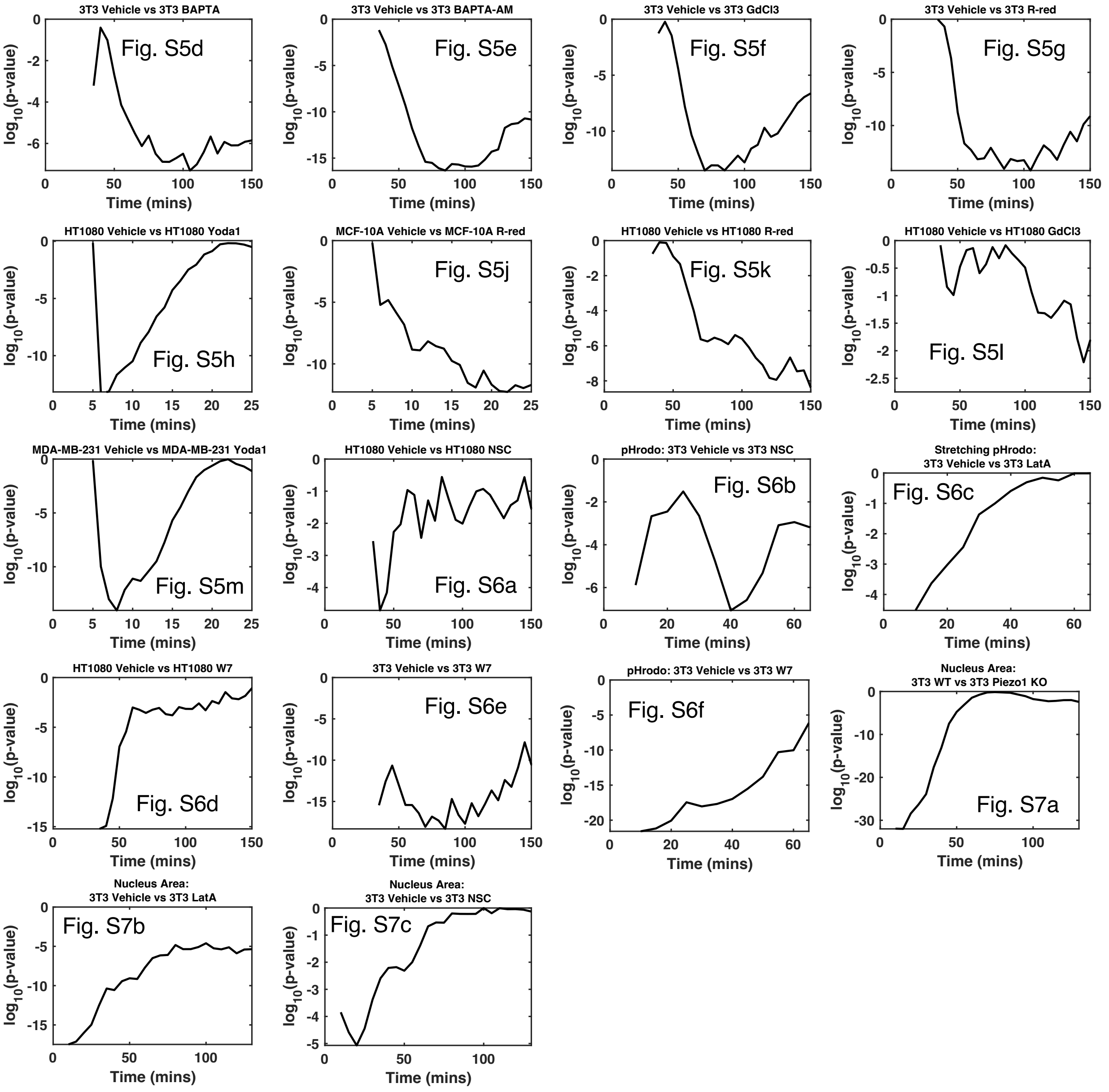


Figure ST-5: Statistical analysis of all time series plots in Supplementary figures S5-S7.

### References


1. Milton H Saier, Vamsee S Reddy, Gabriel Moreno-Hagelsieb, Kevin J Hendargo, Yichi Zhang, Vasu Iddamsetty, Katie Jing Kay Lam, Nuo Tian, Steven Russum, Jianing Wang, and Arturo Medrano-Soto. The Transporter Classification Database (TCDB): 2021 update. *Nucleic Acids Research*, 49(D1):D461–D467, jan 2021.

2. Ruth L Seal, Bryony Braschi, Kristian Gray, Tamsin E M Jones, Susan Tweedie, Liora Haim-Vilmovsky, and Elspeth A Bruford. Genenames.org: the HGNC resources in 2023. *Nucleic acids research*, 51(D1):D1003–D1009, jan 2023.

3. Y. Li, X. Zhou, and S. X. Sun. Hydrogen, bicarbonate, and their associated exchangers in cell volume regulation. *Front. Cell Dev. Biol.*, 9:683686, 2021.

4. H. Jiang and S. X. Sun. Cellular pressure and volume regulation and implications for cell mechanics. *Biophys. J.*, 105(3):609–619, 2013.

5. Y. Li, K. Konstantopoulos, R. Zhao, Y. Mori, and S. X. Sun. The importance of water and hydraulic pressure in cell dynamics. *J. Cell Sci.*, 133(20):jcs240341, 2020.

6. F. Yellin, Y. Li, V. K. A. Sreenivasan, B. Farrell, M. Johny, D. Yue, and S. X. Sun. Electromechanics and volume dynamics in non-excitable tissue cells. *Biophys. J.*, 114:2231–2242, 2018.

7. D. C. Gadsby, J. Kimura, and A. Noma. Voltage dependence of Na/K pump current in isolated heart cells. *Nature*, 315:63–65, 1985.

8. J. R. Casey, S. Grinstein, and J. Orlowski. Sensors and regulators of intracellular pH. *Nat. Rev. Mol. Cell Biol.*, 11(1):50, 2010.

9. A. M. Weinstein. A mathematical model of the rat proximal tubule. *Am. J. Physiol. Renal Physiol.*, 250(5):F860–F873, 1986.
